# Climate change and the complex dynamics of green spruce aphid–spruce plantation interactions

**DOI:** 10.1101/2021.05.26.445772

**Authors:** John H. M. Thornley, Jonathan A. Newman

## Abstract

Aphids can have a significant impact on the growth and commercial yield of spruce plantations. Here we develop a mechanistic deterministic mathematical model for the dynamics of the green spruce aphid (*Elatobium abietum* Walker) growing on Sitka spruce (*Picea sitchensis* (Bong.) Carr.). These grow in a northern British climate in managed plantations, with planting, thinning and a 60-year rotation. Aphid infestation rarely kills the tree but can reduce growth by up to 55%. We used the Edinburgh Forest Model (efm) to simulate spruce tree growth. The aphid sub-model is described in detail in an appendix. The only environmental variable which impacts immediately on aphid dynamics is air temperature which varies diurnally and seasonally. The efm variables that are directly significant for the aphid are leaf area and phloem nitrogen and carbon. Aphid population predictions include dying out, annual, biennual and other complex patterns, including chaos. Predicted impacts on plantation yield of managed forests can be large and variable, as has been observed; they are also much affected by temperature, CO_2_ concentration and other climate variables. However increased CO_2_ concentration appears to ameliorate the severity of the effects of increasing temperatures coupled to worsening aphid infestations on plantation yield.

## 1 Introduction

The impact of climatic change on forests, pest populations, and their interactions, has been the focus of much work over the last several decades. The biology is complex, and involves both direct and indirect effects of multiple climate variables. Elevated CO_2_, changing temperature, and their interactions have direct impacts on plant growth and quality (from the herbivores’ perspectives [1]). Many invertebrate pest species’ population dynamics are highly sensitive to ambient temperature means and variances [2]. While a great deal of empirical experimental work has been done on species of economic consequence, their long-term responses to climatic change can only really be assessed with models. In this paper, we construct a mechanistic ecosystem simulation model of the interaction between Sitka spruce trees (*Picea sitchenis*) and their herbivores the green spruce aphid (*Elatobium abietinum*).

### 1.1 The biological system

The green spruce aphid, *E. abietinum* (syn. *Aphis abietina*), is a significant pest of some species of spruce (*Picea*) in parts of Europe. Its ecology and impacts have been comprehensively reviewed by Day et al. [3]. Dixon [4] gives an excellent introduction to the science of aphids including much material relevant to the green spruce aphid. Sitka spruce (*P. sitchenis*) is an exotic species in northwest Europe; it is now the predominant plantation species in maritime areas, where it produces a yield of 9–15 m^3^ ha^−1^ y^−1^ of stem wood over a rotation — in Scotland with Sitka spruce this is typically 60 years. It is one of the most productive trees in this situation and this is being enhanced by progressive genetic gains (see preface of [3]).

The aphid feeds on phloem sap [5]. It partially defoliates but rarely kills its host; it can depress annual growth by 10–50% [6]. The impact of such infestations is generally summarized by its effect on observed yield of stem wood during and over a rotation. A rotation length of 60 years does not allow studies which directly address the problem to be easily executed. Therefore research remains mostly empirical and short term. Randle and Ludlow ([7] p. 33) state that “The ideal model for defoliation studies remains to be developed” and this seems to remain largely true (but, for studies of aphids in other systems, see Table A.1 for summary).

Interest in this particular tree-aphid system’s response to climatic change dates back to at least the mid-1990s. Straw [8] summarized the assessment at that time as follows (p. 134):

> “Defoliation of Sitka and Norway spruce by the green spruce aphid (*Elatobium abietinum*) is limited in the UK primarily by periods of cold weather which reduce the number of aphids overwintering. If winters become milder, as current models of climate change predict, then the aphid is likely to become more abundant and years with severe defoliation more frequent. In such circumstances the productivity of spruce will decline.”

### 1.2 Models of aphids and climatic change

Probably due to their economic importance, aphids have been the focus of numerous modelling studies. These studies fall into three main types: statistical, agent-based, and mechanistic.

#### 1.2.1 Statistical models

In the context of climatic change, statistical models of insect responses are largely so-called species distribution models (SDMs). Popular tools include Maxent (a form of presence only logistic regression modelling), Genetic Algorithm for Rule-set Production (GARP; [9]) and several others. For a comparsion of these techniques, see [10]. While these methods have been popular for modelling the potential impact of climatic change on non-aphid insects, they have rarely been used for aphids. There are plenty of examples of SDMs for aphids (see e.g. [11–13]) but it is more rare to see these coupled with projections of climatic change (but see e.g. [14, 15]).

Statistical models contain no information beyond the original data used to construct them. They say nothing about the mechanisms that give rise to the response. For example, SDMs struggle to handle species interactions, due to their lack of mechanisms. This can be particularly problematic for herbivore-plant interactions since the host plant responds dynamically to climatic change and ignoring changes in the host plant’s distribution or quality can result in very different views of the future (see e.g. [16, 17]). Also, while it seems reasonable to estimate thermal tolerances from current species distributions, SDMs are incapable of considering changes in CO_2_ concentrations, another key component of climatic change for plant-herbivore interactions [18, 19]. Nevertheless, statistical models are useful for summarizing data and interpolating between data. They are often ‘user friendly’ and can be readily fashioned into tools valuable to farmers or farm advisors.

#### 1.2.2 Agent based models

Another common modelling approach is to use spacially explicit models (SEMs), particularly agent based models. DeAngelis et al. [20] provides a useful review of the approach and its relationship to other modelling approaches. They point out that SEMs can reveal aspects of local and regional level processes that are often absent in spatially implicit models (SIMs). Their review suggests that “spatial models in ecology have largely gone in different directions: towards SEMs for applied or pragmatic problems and towards SIMs for theoretical problems” ([20] pg. 294). This is possibly due to the fact that SEMs require (or at least can make use of) detailed landscape information, the gathering of which can be a long, laborious, and expensive process [20]. Thierry et al. [21] develop a general agricultural landscape modelling framework that can be used to explore the effects of agricultural landscape dynamics on organisms. While agent-based modelling is used widely in ecology, its use for modelling aphid dynamics has been more limited.

Parry et al. [22] constructed an individual-based aphid population model. They modelled a 5 km × 5 km region of Hertfordshire in southeastern England but did not consider climatic change. Agent based models are computationally intensive. As the authors point out, a challenge for this approach is to expand it so that it can cover realistic aphid densities across larger regions, which will increase the run-time and computational power required. Wiest et al. [23] simulated the population growth of the *Rhopalosiphum padi* in wheat plants exposed to environments with different thermal regimes. Population size varied according to the thermal regime. The effects of constant, daily variation, and outside mean minimum/maximum air temperature thermal regimes on the development and fecundity rates were not uniform. Although this model could be used to explicitly study the impacts of climate change, the authors did not do so. Picaza et al. [24] used an agent based model to study aphids as disease vectors, but did not model temperature dependent population growth and so it is not suitable for the study of climate change impacts.

#### 1.2.3 Mechanistic models

There is a long tradition of using mechanistic models to study aphid population dynamics. Table A.1 summarizes a sampling of these models. Commonalities found in these models are readily apparent. First, with the exception of Newman et al. [25–29] these previous models have not been designed for, or used for, studying the impacts of climatic change (but see [30]). Second, since ‘climate change’ encompasses, *at a minimum*, changes in air temperature and CO_2_, none of the previous models are even suited to the task because they do not consider the effects of rising CO_2_ (again, with the exception of the Newman et al. studies). Third, with the exception of Barlow et al. [31–33] and a very basic model by Day et al. [3], none of previous models have considered tree aphids and even these models lack a mechanistic treatment of tree growth and physiology. And fourth, even for models that do include a consideration of the host plant, many of these models do not dynamically link the aphid and the host plant. That is, the models of the plants tend to be very simplified and unresponsive to aphid pressure. On the other hand, Newman et al. [25] constructed a model of cereal aphid population dynamics and coupled it to the Hurley Pasture Model [34, 35], a long-established mechanistic ecosystem simulation model of temperate grass pastures. They [25] used this model to gain insight into the generality of aphid population dynamic responses to climatic change, to understand the magnitude and direction of each of the climate variables’ impacts, and to explore the role of predation in controlling aphid populations under climate change [25, 27–29, 36].

Our objective here is to construct a transparent mechanistic model of the green spruce aphid and interface this with a long-established mechanistic forest ecosystem simulation model, the Edinburgh Forest Model (efm, [37]). This is then used to examine predictions under various climate scenarios. We believe that an understanding of green spruce aphid dynamics can only be obtained by combining a mechanistic aphid model with a mechanistic plant growth model. No tuning (or less charitably - ‘parameter twiddling’) has been applied. At this stage, it is arguably more valuable to examine the range of behaviour the model can predict, than to look for an understanding of the discrepancies which may exist between observation and theory.

## 2 Tree sub-model

The aphid sub-model is interfaced with the Edinburgh Forest Model (efm). The efm is a mature and well-validated mechanistic simulator applicable to evergreen or deciduous forest ecosystems ([37]; see Appendix B below for the numerical methods employed). These can be grown as plantations, managed forests, or unmanaged forests. The model is based on simplified physiology and biochemistry with soil and water sub-models. The efm couples carbon (C), nitrogen (N) and water, fluxes and pools and provides stoichiometric balancing of the items represented. The efm is shown schematically in Fig 1 and sketched in some mathematical detail in Appendix C.

**Fig 1.**
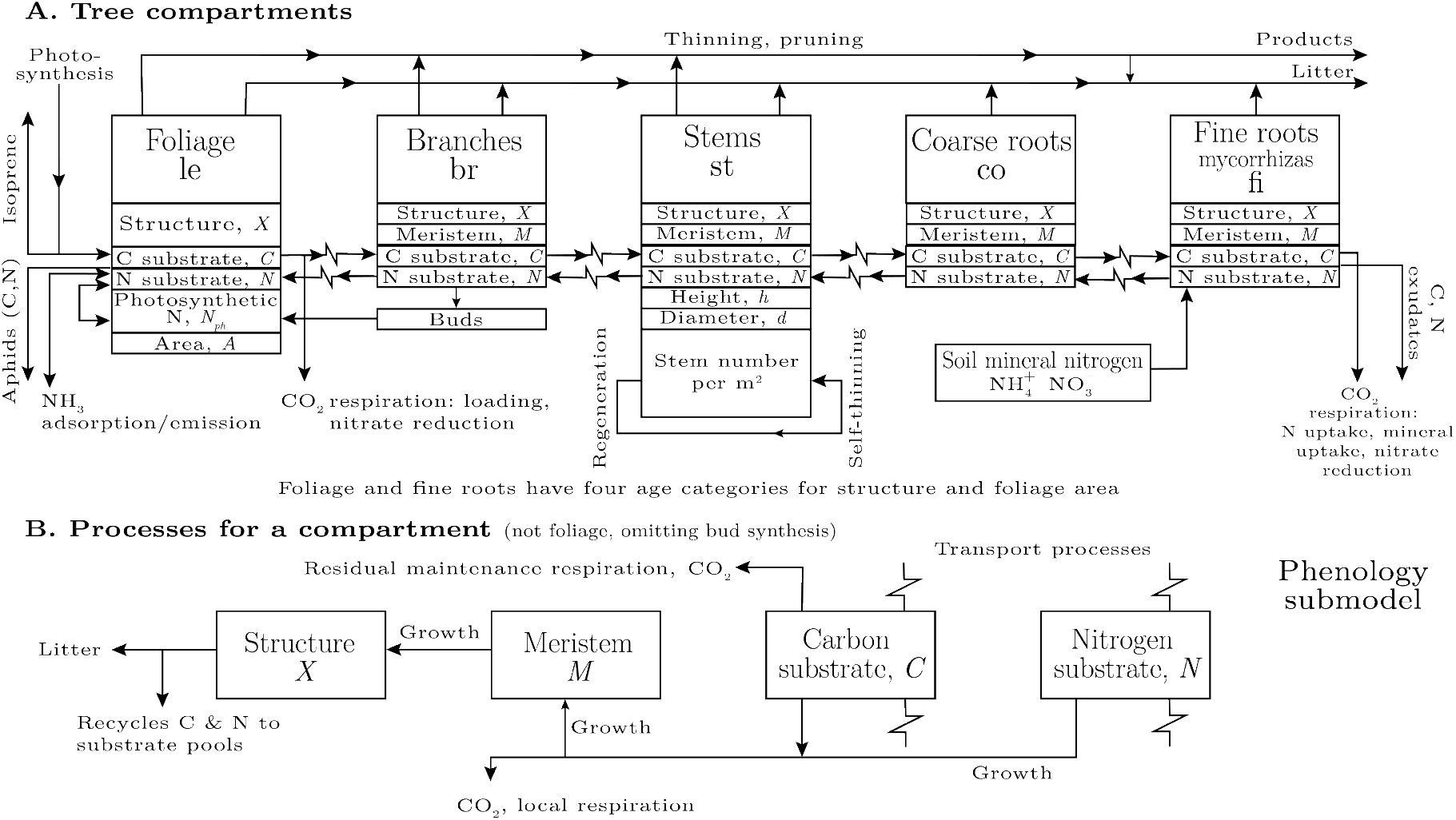
Tree sub-model of the Edinburgh Forest Model (efm). Aphids are connected to the tree sub-model on the left side in A, where aphids extract C and N from the foliage (leaf, le) substrate pools, denoted *C*_le_ and *N*_le_, via the phloem [Eq (2)]. The aphid sub-model can be ‘switched off entirely (the default position).

## 3 Overview of aphid sub-model

In this section we give an overview of the aphid sub-model, which has ten state variables as shown in Fig 2. Details of the ordinary differential equations (ODE’s) which determine their dynamics, the fluxes which contribute to the ODE’s and their behaviour are given in detail in Appendix D.

**Fig 2.**
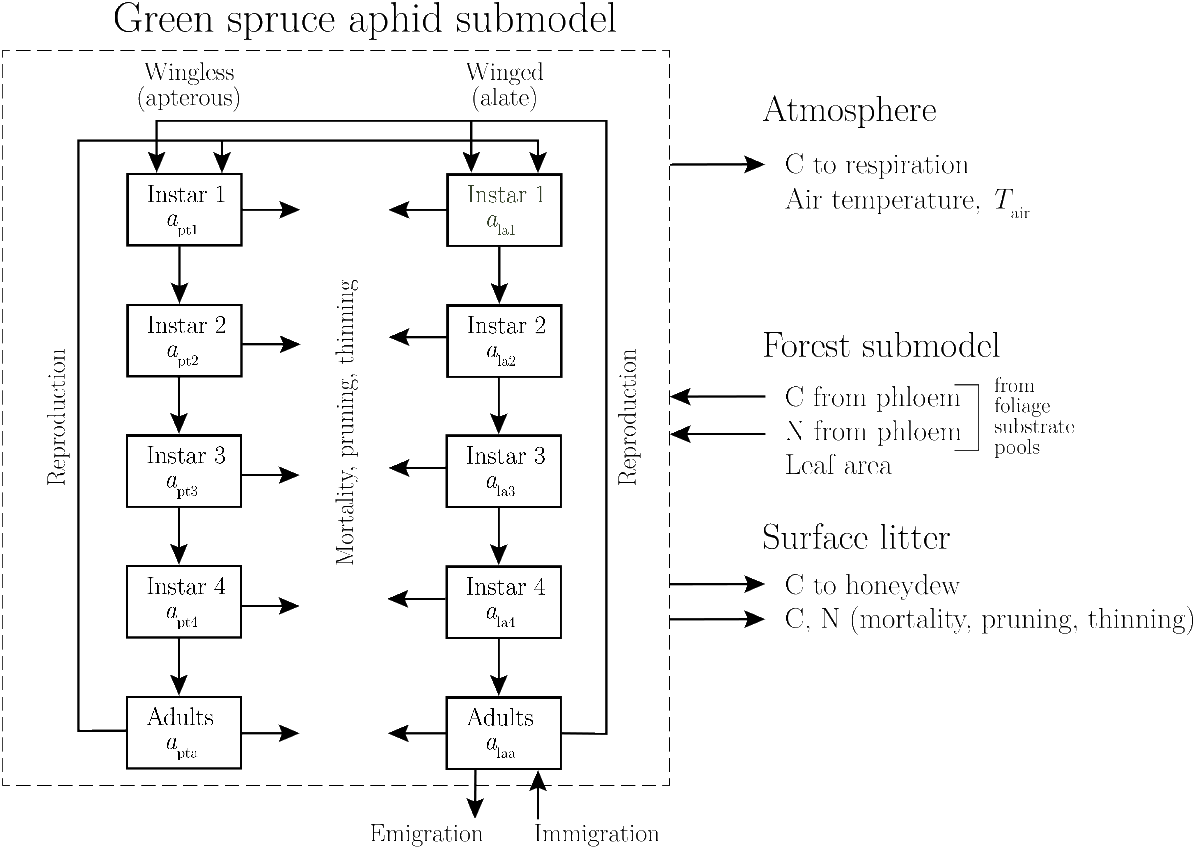
The green spruce aphid sub-model. The ten state variables of the sub-model are shown in the boxes. There are four instars and an adult form for the wingless (apterous) and the winged (alate) aphids. Units for the state variables are number of aphids per stem (Appendix D). The differential equations for the state variables are Eqs (16), (47), (64) and (87).

## 4 Simulation scenarios

### 4.1 The environment

To illustrate the model’s dynamics, we used the average climate for Eskdalemuir, in northern Britain, latitude 55°19′ N, longitude 3°12′ W, altitude 242 m above sea level (Fig 3). The monthly data [38] were supplemented by monthly data from Clino [39]. Daily values were obtained by linear interpolation of monthly data (see chapter 7 and Section 7.5.3 of Thornley [34]). Daily data were applied to give diurnally changing data for air temperature (Fig 3A), relative humidity (Fig 3A), radiation (which is calculated from sunshine hours; Fig 3B), and wind (not shown). Soil temperature and rainfall were assumed constant over each day. The source program, `efm.csl`, gives details (see Appendix C).

**Fig 3.**
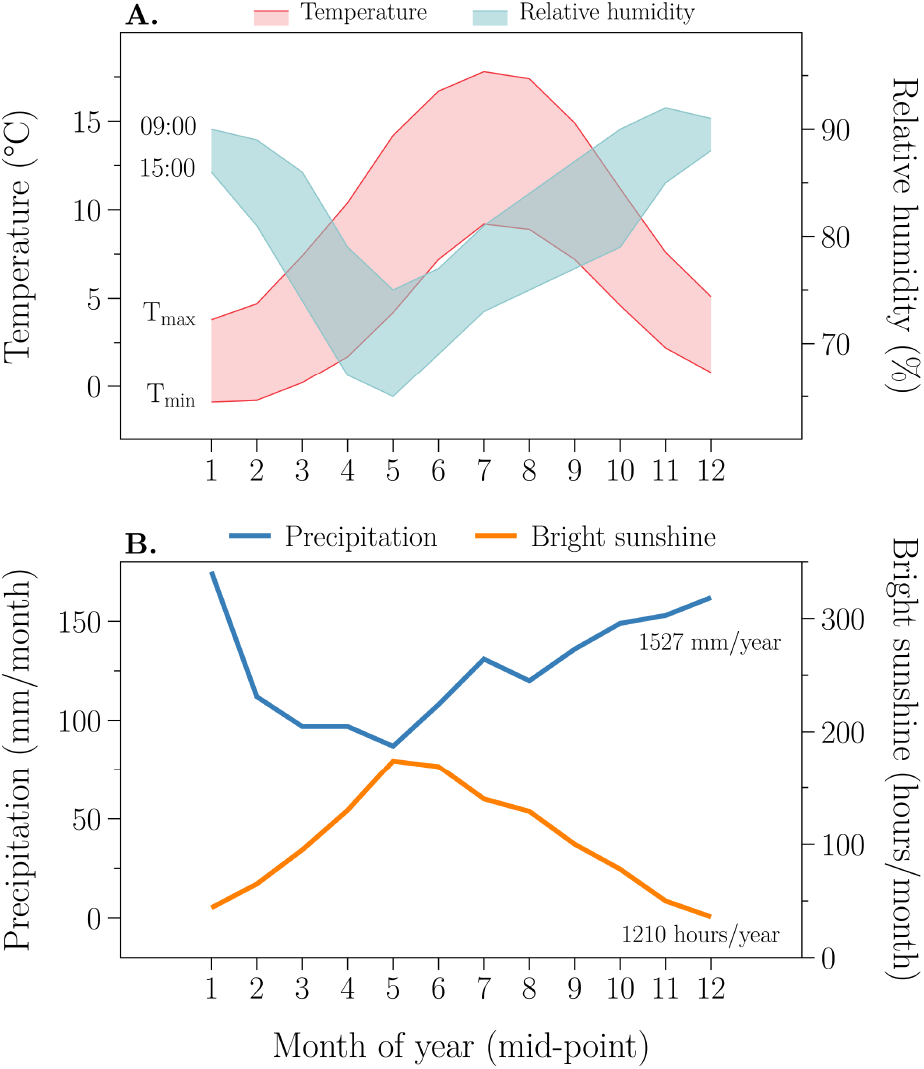
Climate at Eskdalemuir. 30-year monthly means taken from meteorological tables are drawn. See text for details.

Wind speed at 50 m reference height (*h*_ref_) varies seasonally with the maxima of the daily maxima and daily minima occurring on 26 April. The mean and the variation of the daily maximum wind speeds are 6 and 1.5 m s^−1^. The mean and the variation of the daily minimum wind speeds are 2 and 0.5 m s^−1^. The mean annual wind speed is 1/2 × (6 + 2) = 4 m s^−1^. We assumed that the diurnal variation of wind speed is similar to that of air temperature (*T*_air_) and opposite to that of relative humidity (RH): the minimum wind speed (and minimum temperature and maximum RH) occurs at dawn and the maximum wind speed (and maximum temperature and minimum RH) at 15 h.

Photosynthetically active radiation (PAR) varies from a minimum of 0.55 on 20 December to a maximum of 7.9 MJ PAR m^−2^ d^−1^ on 15 June. It is calculated from bright sunshine hours (Fig 3B) by using a version of the Ångström formula ([34], pp. 145–146, equation 7.4i, but with *a*_Ang_ = 0.19 and *b*_Ang_ = 0.62; [40]) for daily PAR light receipt, *j*_PARdy_ (J m^−2^ d^−1^) from the fraction of bright sunshine hours which has been interpolated from monthly values (Fig 3B) to give daily values and refers to a given Julian day number.

Soil temperature was assumed to be diurnally constant and equal to mean daily air temperature; it varies from a minimum of 1.45 °C on 16 January to a maximum of 13.5 °C on 16 July. Diurnal air temperature variation is a maximum of 5 °C on 16 May and a minimum of 2.15 °C on 16 December.

Relative humidity (RH) was assumed to be at its daily maximum at dawn and at its daily minimum at 15 h. The RH daily maximum has a minimum of 0.75 on 16 May when the RH daily minimum is 0.65 and the RH daily minimum has a maximum of 0.88 on 16 December when the RH daily maximum is 0.91.

Daily rain fall (assumed to be diurnally constant) is substantial, varying from a minimum of 2.8 kg (2.8 mm) m^−2^ d^−1^ on 16 May, to a maximum of 5.65 kg (5.65 mm) m^−2^ d^−1^ on 16 January, with an annual rain fall of 1527 kg (1527 mm) m^−2^ y^−1^ (see Fig 3A).

### 4.2 Forest plantation

A regime of planting to an initial stem density of 0.25 stems m^−2^ and an initial leaf area index of 0.003 [Eq (1)], followed by the removal of 0.45, 0.4, 0.35, 0.3, 0.25, 0.2, 0.15 and 0.1 of the existing stems at times of 20, 25, 30, 35, 40, 45, 50 and 55 years and terminated by clear felling at 60 years, was employed. Stem removal takes place at a constant rate over a period of 1 d (1 January normally) and this determines the thinning function *O*_nstems,th_ of Eq (1), which depends on the integration step, maxt (Appendix B).

### 4.3 Climate change simulations: temperature and CO_2_

Various temperatures (−3 to +4 °C) below and above the ambient Eskdalemuir temperatures and two CO_2_ concentrations: 350, 700 μmol mol^−1^ were applied. Although current ambient CO_2_ is now above 400 μmol mol^−1^, we chose 350 μmol mol^−1^ as “current ambient” to aide comparison with previous experimental and theoretical research. Initial values for all efm state variables were equilibrium with no aphids for the temperature and CO_2_ level applied. The equilibrium is a 60-year repeating equilibrium; the same 60-year rotation period is applied to all the temperature × CO_2_ scenarios considered, although this would not usually give the optimum timber yield (yield class, *Y*_C_) for all cases.

## 5 Results

### 5.1 Aphid population dynamics

In Fig 4 the aphid population densities (ρ_aph_) are plotted together, 700 vpm CO_2_ (red) overlaid on 350 vpm CO_2_ (black), for each incremental temperature change (Δ*T*) from −2 to +3 °C for the full 60 year rotation. Also shown are the corresponding spectral densities^1^ calculated for the final 32 years of the rotation. Peaks in the spectral density plots indicate periodicity [41]. Fig 4 demonstrates *qualitative differences* in the population dynamics. At −2 °C, aphids more or less die out well before the 60 year rotation is complete (they are still present, but at vanishingly low densities), and the die out happens faster under 700 vpm CO_2_. At −1 °C, both CO_2_ conditions transition to essentially the same, reasonably stable, annual cycles of relatively low aphid densities (ρ_aph_), although they do so on different trajectories and the spectral density shows that there is additional underlying within year structure. At ambient temperature (Δ*T* = 0), both CO_2_ conditions begin with annual cycles but diverge and transition to chaotic dynamics. The corresponding spectral densities have no obvious structure, indicative of (though not determinative of) chaos. At +1 °C, aphids transition from annual cycles to more complex periodicities. The spectral densities show that the periodicities are different and out of phase with each other. At +2 °C we see a similar pattern, but with stronger signals of periodicity. Finally, at +3 °C and 350 vpm CO_2_, aphids again show sustained 2-point cycles, while at 700 vpm CO_2_ the annual cycles are supplemented with complex within year dynamics.

**Fig 4.**
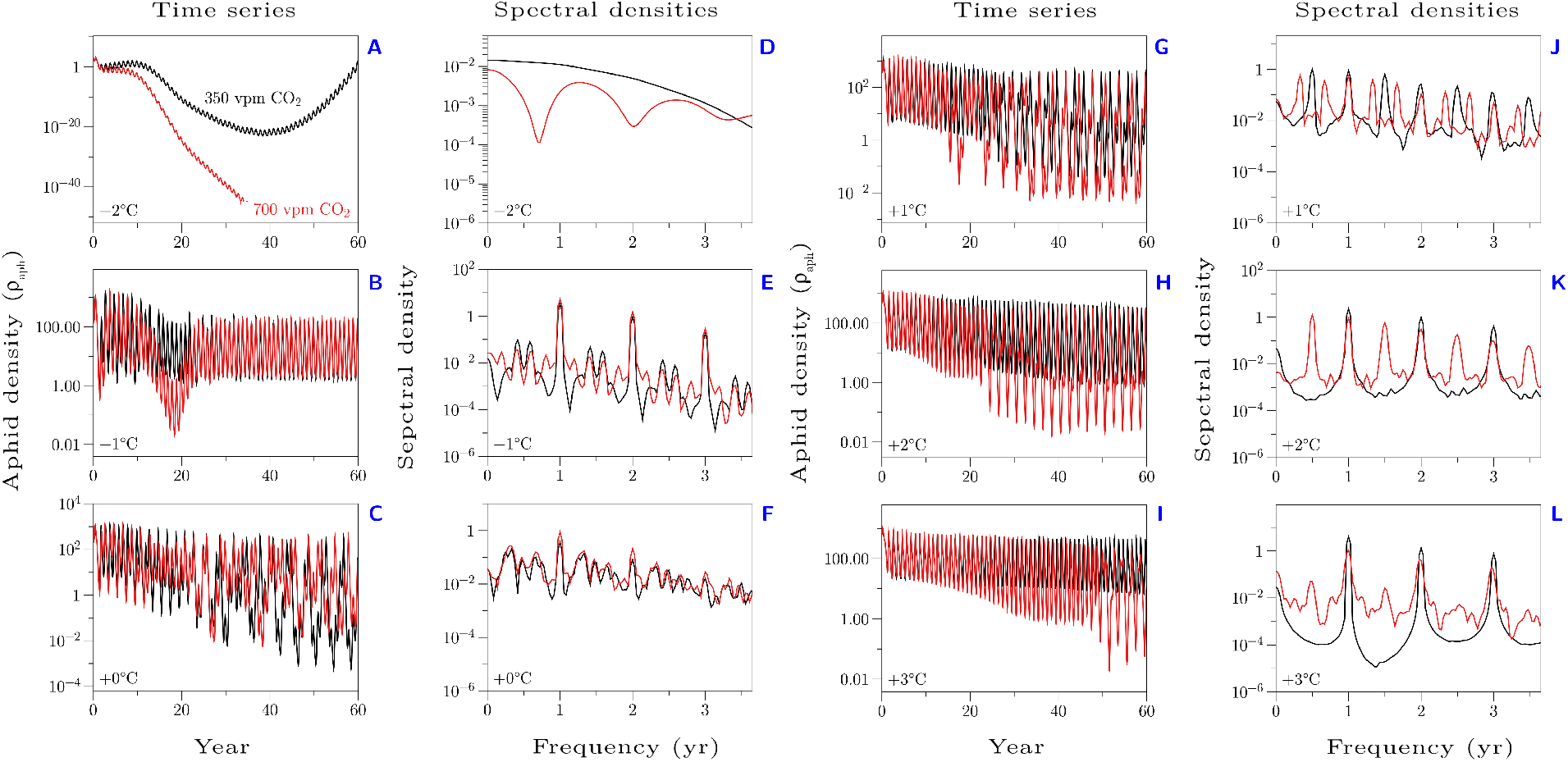
Aphid density dynamics. Shown are the time series of aphid densities (ρ_aph_) for incremental increases in temperature from —2 °C to +3 °C. 350 vpm CO_2_ is shown in black, and 700 vpm CO_2_ is shown in red. Notice the qualitative differences in population dynamics that emerge as temperture and CO_2_ change. Next to each time series is the respective spectral density (arbitrary units) calculated from the last 32 years of each time series; the frequency (*x*-axis) has been re-scaled to display in years.

We can compare the dynamics depicted in Fig 4 to those seen in Fig 5. Here we plot the normalized trap captures of alate *E. abietinum* using data extracted from Day et al. [42]. We can see that this time series too contains periodic structure verging on chaos. The spectral decomposition has less structure than we see in our simulations. While we show the spectral decomposition for the same length of time (32 years) our ‘sampling’ of that period is 73 times more dense. Nevertheless, Fig 5 shows one feature that is also seen in our simulations, within year cycles (see e.g., Fig 4 +1 °C, either vpm CO_2_ concentration).

**Fig 5.**
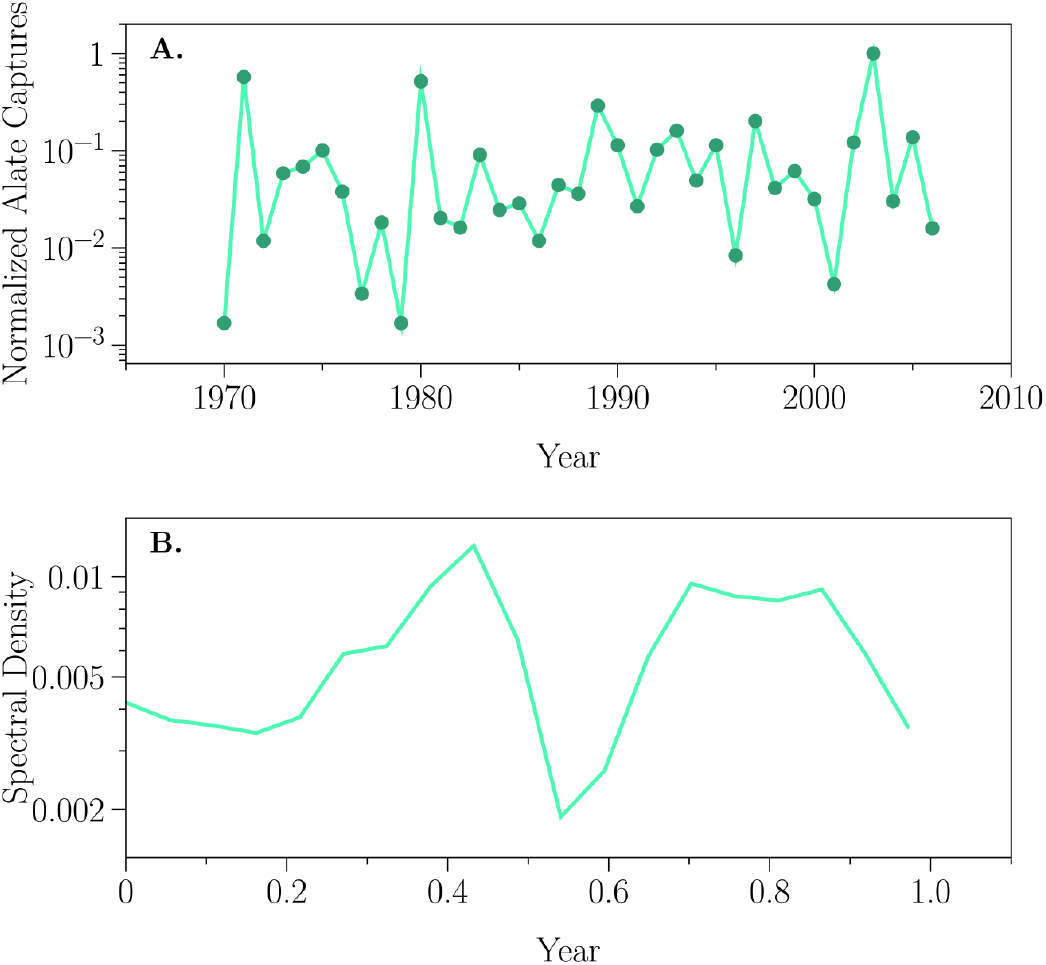
Alate density dynamics. Shown are the time series of alate aphid captures (normalized to the maximum number of captures, see Day et al. [42] for more details) and the associated spectral density function.

### 5.2 Leaf Area Index (*L*_AI_)

Fig 6 shows the Leaf Area Indices that result from the aphid density dynamics. *Without* aphids at 350 vpm CO_2_, increasing temperature generally increases the resulting LAIs, although toward the end of the 60 year rotation the ambient temperature ends up resulting in the highest LAI. Doubling the CO_2_ results in greater values of LAI for all temperature increments, although by the end of the rotation the different temperature trajectorys largely converge. *With* aphids, we see striking differences. At 350 vpm CO_2_, we see a complete reversal in the ordering of the LAIs. Now, −2 °C results in the greatest LAI because aphid population densities are the lowest in these conditions (Fig 4A). A doubling of CO_2_ while generally resulting in higher LAIs does not fundamentally change the conclusion. −2 °C still results in the greatest LAI, because aphid densities are still lowest at this temperature (Fig 4A).

**Fig 6.**
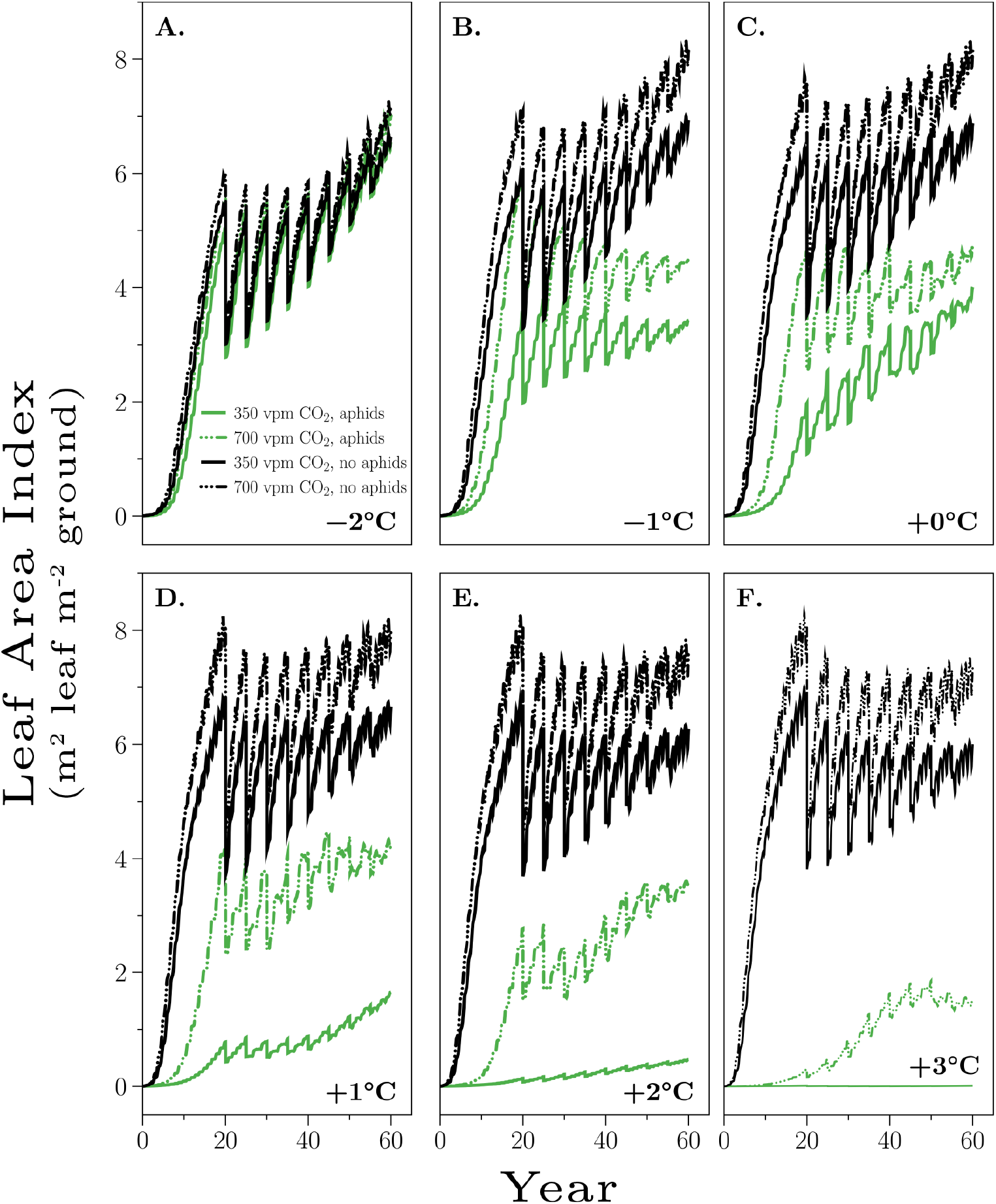
Leaf Area Indices. Shown are the Leaf Area Indices (m^2^ leaf m^−2^ of ground), Eq (1), for incremental temperature changes at 350 vpm and 700 vpm CO_2_, with and without aphids. The LAI values result from changes in aphid density dynamics, see Fig 4.

### 5.3 C-sequestration (*C*_sys_) and plantation yield (*Y*_C_)

The carbon-sequestration (*C*_sys_) and plantation yields (*Y*_C_) follow from the effects of aphids density dynamics, and temperature and CO_2_ concentration changes. Fig 7 shows the four combinations of aphid presence and CO_2_ for each temperature increment. It is clear from the figure that doubling CO_2_ results in greater C-sequestration and greater plantation yields at temperatures of −1 °C to +4 °C. The presence of aphids results in lower C-sequestration and lower plantation yields. In general these two effects seem to be approximately additive.

**Fig 7.**
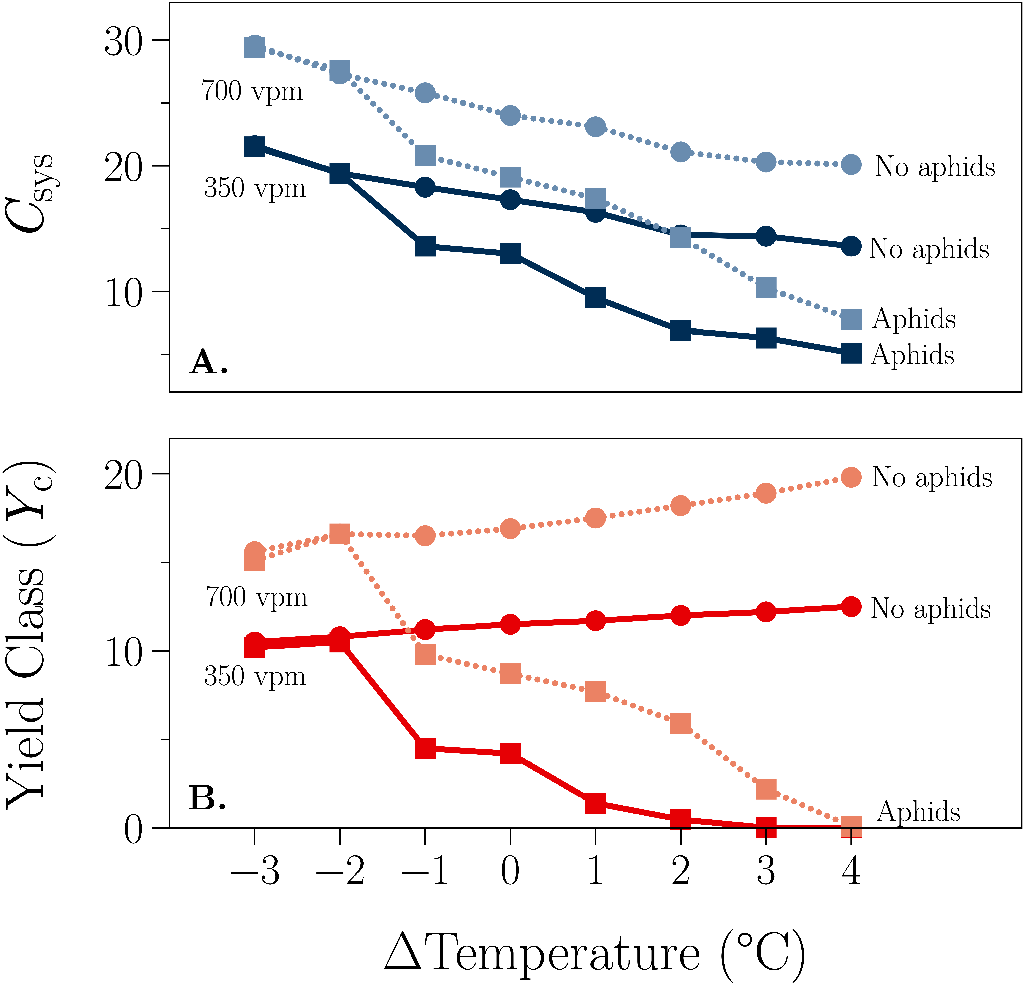
Yield and C-sequestration. ΔTemperature is the increment in air and soil temperatures applied to the Eskdalemuir environment (Section 4.1, Fig 3). The CO_2_ concentration is denoted by the shading (dark color: 350 vpm, light color: 700 vpm). The aphid condition is denoted by the symbol (●: without aphids, ■: with aphids). *Y*_C_ is the yield class (m^3^ ha^−1^ y^−1^), defined as the volume (m^3^) of timber harvested at the end of a rotation per hectare averaged over the duration of a rotation of 60 years; *C*_sys_ (kg Cm^−2^) is the total C in the system at the end of the 60 y rotation. Results are shown for the no-aphid infestation situation (a steady state as applied in Fig 4) and for that when aphids are applied at time zero [Eq (64)].

## 6 Discussion and conclusions

The dynamics of the aphid populations is interesting due to its variety, ranging from aphids dying out (Fig 4A, −2 °C) to various degrees of chaos with annual, biennial and triennial cycles (Fig 4). It is not surprising that chaos occurs rather robustly in the aphid-plant context. There is much non-linearity prevalent; there are far more than the minimum two state variables required for chaos; the equations are non-autonomous; and the intrinsic time scales of the dynamics of the aphid sub-model and the tree sub-model are very different and incommensurate [43, 44]. This chaotic variety indicates that focusing on management/control strategies in order to minimize adverse consequences may be difficult, even with the assistance of a process-based model (for further consideration of chaotic aphid population dynamics see [45–49]).

### 6.1 Comparisons with modeling studies

The model described here is far from the first model to describe the population dynamics of aphids using simulation modelling. Appendix A summarizes 40 aphid simulation modelling studies. Where the present model differs from all but Newman et al. [25–29] is in (*a*) the treatment (mechanistic or otherwise) of the impacts of rising CO_2_ on plant growth and (*b*) the use of the model to study impacts of climatic change. Both the present study and Newman et al. found that the mechanistic consequences of rising temperature tends to benefit aphid population growth while rising CO_2_ tends to be detrimental due to the changes in the plant caused by the rising CO_2_ (see also [26]).

Although there are similarities between the present model and that of Newman et al. [25], they differ in several important respects. For example, Newman et al.’s model is not stochiometrically complete as is the current model. Newman et al. also used their model to answer different questions, focused almost exclusively on the question of changes in aphid abundance. However, perhaps the most significant difference between the two studies is that Newman et al. use their model to study *within* year dynamics, while we used the present model to study an entire 60-year plantation rotation. By studying the dynamics over such a long period of time, we were able to predict the responses illustrated in Fig 4, which range from aphids dying out at low temperatures, to aphids having a severe impact on the productivity of managed spruce plantations (Fig 7B). Our work suggests that this may be a promising approach to investigating the impact of aphids on plant ecosystems in a changing climate. Our simulations indicate that, while temperature is the most important environmental variable, higher CO_2_ levels ameliorate the impact of aphid infestation on yield and carbon sequestration (Fig 7) and on leaf area index (Fig 6) and aphid dynamics (Fig 4). None of the previous modelling exercises have been able to provide such an integrated picture of the possible impacts of climatic change, let alone for plantation ecosystems.

### 6.2 Model criticism, limitations and extensions

The aphid model is so far untuned for any specific purpose although where possible parameter values were estimated for the green spruce aphid. Our analysis leans heavily on Dixon [4] and on Day et al. [3] who focus specifically on the green spruce aphid. While we have tried to tie parameter values down to data and predictions to observations, this has been difficult, perhaps due to the traditions in this area of work. As mentioned earlier, we have not done any indiscriminate ‘parameter-twiddling’, which can be an endless process. With any model, it is preferable to fix parameters by experiments targeted at the level of the parameters than by adjustment with reference to the outcomes predicted by the model which depend on everything within the model; we know that all models are wrong in some respect. We have found that the occurrence of chaos and our general results are qualitatively not greatly affected by the details of parameterization, so long as the values assumed are in the right ball-park. The main conclusion is, we suggest, that mechanistic aphid-plant models may be an essential but difficult approach to understanding how these systems work.

## 7 Acknowledgments

The representation and parameterization of the Edinburgh Forest Model owe much to a long collaboration with Melvin Cannell and colleagues at the Institute for Terrestrial Ecology, Penicuik, Edinburgh. The work reported here is partly supported by the UK Department for Environment and Regions, contracts 1/1/160 and 1/1/64. JAN was supported by grants from the Canadian Natural Science and Engineering Research Council and from the Ontario Ministry of Agriculture, Food, and Rural Affairs. Wilfrid Laurier University and its campuses are located on the Haldimand tract, traditional territory of the Neutral, Anishnaabe and Haudenosaunee peoples. We recognize, honour, and respect these nations as the traditional stewards of the lands and water on which Laurier is now present.

## Appendix A Aphid simulation models

**Tab A.1.** Summary of aphid simulation models. ‘TD aphid’ denotes that aphid growth is represented in the model as temperature-dependent; similarly ‘TD plant’ denotes that plant growth is represented as temperature-dependent. ‘CO_2_ plant’ denotes whether the model considers the CO_2_ concentration and its mechanistic effects on plant growth. ‘Climate Δ’ denotes whether climate change is explicitly considered or not. The ‘Stated purpose’ indicates the purpose of the model according to the authors. ‘??’ denotes that the information was not presented in the noted reference.

| Reference (Year) | TD aphid | Plant species | TD plant | CO<sub>2</sub> plant | Climate Δ | Stated purpose |
| --- | --- | --- | --- | --- | --- | --- |
| Aphids that feed on graminoid host plants |  |  |  |  |  |  |
| Bird cherry-oat aphid (*Rhopalosiphum padi* L.) |  |  |  |  |  |  |
| Wikteliuss & Pettersson [50] (1985) | Yes | None | N/A | N/A | No | “A model of the population dynamics of the bird cherry-oat aphid (*Rhopalosiphum padi* (L.)) has been used to study the likely effects of plant resistance on aphid population growth.” |
| Ekbom et al. [51] (1992) | Yes | Spring barley | Yes | No | No | “The models were used to answer several questions of importance for evaluation of naturally occurring control. 1) Can polyphagous predators find aphids at very low prey densities? 2) Is the timing of predation in relation to the aphid’s phenology important for resulting population development? 3) What are the necessary conditions for effective control by natural enemies?” |
| Morgan [52] (2000) | Yes | Winter barley | No | No | No | “The aims of this paper are to utilize simulation analysis techniques to develop a model describing the population development of *R. padi*, to use the model to examine the relative importance of various population processes in pest dynamics and to determine whether the model could be used as a basis for forecasting aphid outbreaks.” |
| Newman et al. [25–29] (2003–2006) | Yes | Grass | Yes | Yes | Yes | “In this paper, we develop a mechanistic mathematical model of the general interaction between grasses, cereal aphids and their parasitoids. We used this model to investigate the interacting effects of rising CO<sub>2</sub> and temperature.” |
| Parry et al. [53] (2006) | Yes | None | N/A | N/A | No | “[Used] to investigate the effects of different pesticide regimes.” |
| Thackray et al. [54] (2009) | No | Wheat | No | No | No | “BYDV PREDICTOR, a simulation model, was developed to forecast aphid outbreaks and Barley yellow dwarf virus (BYDV) epidemics in wheat crops in the grainbelt region of southwest Australia.” |
| Russian wheat aphid (*Diuraphis noxia* Kurdjumov) |  |  |  |  |  |  |
| Ma et al. [55] (2008) | Yes | Barley | Yes | No | No | None stated |
| continued on next page |  |  |  |  |  |  |

| continued from previous page |  |  |  |  |  |  |
| --- | --- | --- | --- | --- | --- | --- |
| Reference (Year) | TD aphid | Plant species | TD plant | CO<sub>2</sub> plant | Climate Δ | Stated purpose |
| English Grain Aphid (*Sitobion avenae* Fabricius) |  |  |  |  |  |  |
| Carter et al. [56, 57] (1980) | Yes | Winter wheat | Yes | No | No | “[Used] to explain the population development of *S. avenae* on cereals and to indicate how cereal aphid outbreaks might be predicted.” |
| Rabbinge et al. [58, 59] (1979) | Yes | Wheat | Yes | No | No | “[D]eveloped simulation models that explain the population movements of the most important aphid *S. avenae* and may be used for extrapolation and prognosis in the short run, approximately 3-5 weeks.” |
| Skirvin et al. [30] (1997) | Yes | None | N/A | N/A | Yes | “This paper describes a mechanistic computer model of the interaction between the cereal aphid *Sitobion avenae*, ... and its main predator in Britain, the coccinellid *Coccinella septempunctata*.” |
| Rossing [60] (1991) | Yes | Winter wheat | Yes | No | No | “To evaluate the relative importance of various components of damage caused by grain aphid (*Sitobion avenae* E) populations in winter wheat, a simulation model of crop growth and development is combined with a model of aphid injury.” |
| Freier et al. [61, 62] (1996, 2001) | Yes | Winter wheat | Yes | No | No | “[Used] to study and demonstrate tri-trophic interactions, particularly aphid population dynamics influenced by antagonists under different natural and man-made conditions, but not to predict future aphid infestation.” |
| Duffy et al. [63] (2017) | Yes | Cereal crop | No | No | No | “[N]ew simulation model that quantifies the size and timing of grain aphid populations in response to temperature.” |
| Rossing [60] (1991) | Yes | Winter wheat | Yes | No | No | “A dynamic, process-based simulation model is used to evaluate the effect of injury by *Sitobion avenae* on yield of winter wheat at a range of attainable yield levels.” |
| van Roermund et al. [64] (1986) | Yes | Winter wheat | Yes | No | No | “To study the effect of the English grain aphid, *Sitobion avenae* (F.), on dry matter accumulation in winter wheat grains, the influence of aphids on plant physiological parameters was introduced into a simulation model of wheat growth and development from anthesis onwards.” |
| Aphids that feed on forb host plants |  |  |  |  |  |  |
| cabbage aphid (*Brevicoryne brassicae* L.) |  |  |  |  |  |  |
| Hughes & Gilbert [65] (1968) | Yes | None | N/A | N/A | No | “[T]ool to understand the combined effects of species interactions and environmental factors on population growth |
| Gilbert & Hughes [66] (1971) | Yes | None | N/A | N/A | No | “... refinement used to examine biocontrol strategies.” |
| Raworth [67] (1984) | Yes | kale | Yes | No | No | “This paper consolidates information from the previous studies in the form of a simulation model.” |
| continued on next page |  |  |  |  |  |  |
| Reference<br>(Year) | TD<br>aphid | Plant<br>species | TD<br>plant | CO<sub>2</sub><br>plant | Climate<br>$\Delta$ | Stated purpose |
| pea aphid (*Acyrtosiphon pisum* Harris) |  |  |  |  |  |  |
| Gutierrez et al. [68] (1984) | Yes | Alfalfa | ?? | No | No | “[E]xpands a model for energy or biomass assimilation by individuals, that has been proposed in various forms in the literature, to model the growth, development, reproduction, ageing and survivorship of the pea aphid.” |
| Snyder & Ives [69] (2003) | No | None | N/A | N/A | No | “We investigated how specialist and generalist natural enemies differ as biological control agents of pea aphids (*Acyrtosiphon pisum*), and how interactions among natural enemies affect successful control.” |
| black bean aphid (*Aphis fabae*) |  |  |  |  |  |  |
| Kindlmann [70] (1985) | No | None | N/A | N/A | No | not stated |
| cotton aphid (*Aphis gossypii*) |  |  |  |  |  |  |
| Xia et al. [71] (2018) | Yes | None | N/A | N/A | No | “To better understand and characterize the predator–prey system dynamics under field conditions.” |
| cowpea aphid (*Aphis craccivora* Koch) |  |  |  |  |  |  |
| Gutierrez et al. [72,73] (1974) |  | Clover | Yes | No | No | “... describing the interactions of various components of the biology of cowpea aphid (*Aphis craccivora* Koch), its host plant, Subterranean Clover Stunt Virus (SCSV).” |
| green peach aphid (*Myzus persicae* Sulz.) |  |  |  |  |  |  |
| Whalon & Smilowitz [74] (1979) | Yes | Potato | No | No | No | “This is a report of predictive temperature-dependent density equations for green peach aphid field populations.” |
| Tamaki & Long [75] (1978) | Yes | None | N/A | N/A | No | “A model to evaluate potential reductive impact of populations of green peach aphid, *Myzus persicae* (Sulzer), by predators was generalized to include effects of temperature on the rate of aphid reproduction and the functional response of the predator complex.” |
| Ro & Long [76] (1999) | Yes | Potato | No | No | No | “[T]he model describes the interaction among the green peach aphid population, its predators, and abiotic factors (e.g. temperature). The model includes functions for population growth of the aphid and its predator complex.” |
| continued on next page |  |  |  |  |  |  |
| Reference (Year) | TD aphid | Plant species | TD plant | CO<sub>2</sub> plant | Climate $\Delta$ | Stated purpose |
| potato aphid (*Macrosiphum euphorbiae* Thomas) |  |  |  |  |  |  |
| Bertschinger et al. [77] (1995) | Yes | Potato | Yes | No | No | “A model (EPIVIT) was developed for the simulation of potato harvest infection (% infected tubers) with a contact- or aphid-transmitted virus.” |
| soybean aphid (*Aphis glycines* Matsumura) |  |  |  |  |  |  |
| Miksaneck & Heimpel [78] (2019) | Yes | None | N/A | N/A | No | “We apply this approach to investigate the effect of parasitism by the Asian parasitoid *Aphelinus certus* on its host, the soybean aphid (*Aphis glycines*).” |
| Aphids that feed on tree host plants |  |  |  |  |  |  |
| green spruce aphid (*Elatobium abietinum* Walker) |  |  |  |  |  |  |
| Day et al. [42] (2010) | Yes | None | N/A | N/A | No | “[T]he patterns of insect abundance on trees have never been satisfactorily described by a numerical model despite considerable knowledge of endogenous and exogenous factors in the population dynamics of the species.” |
| lime aphid (*Eucallipterus tiliae* L.) |  |  |  |  |  |  |
| Barlow [31–33] (1979–1981) | Yes | Lime tree | No | No | No | “This paper considers two approaches to the modelling of aphid populations. Using the lime aphid (*Eucallipterus tiliae* L.) as an example, it discusses the well-established role of detailed simulation models in the study of aphid population dynamics then considers the possible application of a simple, general herbivore/plant model to aphids.” |
| Aphids that feed on vine host plants |  |  |  |  |  |  |
| thimbleberry aphid (*Masonaphis maxima* Mason) |  |  |  |  |  |  |
| Gilbert & Gutierrez [79] (1973) | ?? | Thimbleberry | ?? | ?? | No | “[To] explore effects of parasitism and competition on aphid population growth.” |

## Appendix B Numerical methods

The efm is programmed in ACSL, Advanced Continuous Simulation Language, a highly readable fortran-like language (Aegis Research Corporation, Huntsville, AL -) and is extensively commented. ACSL follows the long established CSSL (Continuous System Simulation Language) standard ([80])—designed for models which can be represented by ordinary differential equations using the ‘rate-state’ formalism (rate of change is a function of state variables + driving variables + parameters). A significant advantage of ACSL is that it is non-procedural—that is, the statements can be placed in any order; in the first step, translation, places the statements in an executable order, which can then be compiled and linked as usual. The non-procedural capability allows the programmer to put the statements in an order that makes biological sense. The version of ACSL used here is 11.8.4—now unfortunately legacy software, although to our knowledge there is no adequate replacement. We have used double precision (single precision fails), with translator and runtime table sizes of 200000. The compiler used is Compaq Visual Fortran 6.6-1877-47BAQ, compiling in 36 s. The linker is Microsoft (R) Incremental Linker Version 6.00.8447. Those wishing to use this excellent and much used software system, may contact Professor Parsons at Massey University, New Zealand). The program, efm.csl, which includes the aphid sub-model as an option, is freely available from the senior author. It can also be downloaded from https://sites.massey.ac.nz/hurleypasturemodel/edinburgh-forestry-model/. The time step (maxt) is about 11 minutes (1/128 d) and Euler’s method of integration is applied ([81], pp. 27-40). It takes about eight seconds to simulate one year on a 1.6 GHz pc running under Windows 7 with a 32-bit operating system.

## Appendix C Edinburgh Forest Model

We couple the aphid sub-model (see next section) with the Edinburgh Forest Model (efm). The efm is a mature and well-validated mechanistic simulator applicable to evergreen or deciduous forest ecosystems ([37]. See Appendix B for the numerical methods employed). These can be grown as plantations, managed forests, or unmanaged forests. The model is based on simplified physiology and biochemistry with soil and water sub-models. The efm couples carbon (C), nitrogen (N) and water, fluxes and pools and provides stoichiometric balancing of the items represented.

Here we used the efm in the evergreen forest mode, parameterized for Sitka spruce growing in a north British environment (Section 4.1). The elements of the efm relevant to the aphids are described here. In evergreen mode, foliage is always present and the aphids can live on the foliage year-round, with (in some situations) negligible immigration and emigration. Horizontal homogeneity is assumed. The efm comprises linked sub-models which are described elsewhere for the trees [82], soil and litter ([34], chapter 5) and water ([83]; [34], chapter 6). Synopses of processes represented are given by Thornley [37], Thornley and Cannell [84] and Cannell et al. [85]. Recent developments of the model are: acclimation of photosynthesis to light, nitrogen, carbon dioxide and temperature [86]; components of respiration are more explicitly itemised [87]. See also the appendix of Thornley and Cannell [88] for diagrams of the efm.

Symbols relevant to the present application are listed in Appendix E. Important tree variables vis-à-vis the aphid sub-model are the leaf area per stem, *A*_leaf_ (m^2^ stem^−1^); stem density, *n*_stems_ (number of stems per m^2^, a state variable of the efm); leaf area index, LAI [m^2^ leaf (m^2^ ground)^−1^]; these are given by:

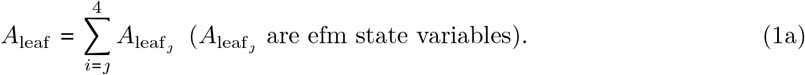

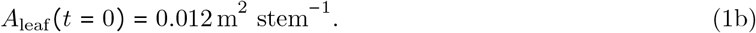

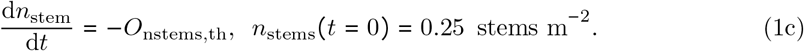

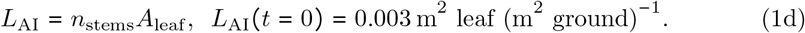

In Eq (1a), leaf area per stem, *A*_leaf_, is represented by four age categories, 1–4 (Fig 1). The time *t* = 0 values each of these variables is 0.003 m^2^ stem^−1^. In the forest plantation simulations (Section 4.2) aphid infestation is assumed to occur at *t* = 0 [Eqs (4), (5) and (64)].

In the second equation, there is a single flux out of the *n*_stems_ state variable, *O*_nstems,th_ (stems m^−2^ d^−1^) due to the management thinning regime (Section 4.2, first paragraph). Pruning is not included in our present simulations of aphid infestation; it is included in the model but is set to zero [Eq (14)].

Other important variables for the aphid implementation of the efm are the carbon (C) substrate and nitrogen (N) substrate concentrations in the foliage (leaf, le), *C*_le_ and *N*_le_. Although the phloem C and N concentrations are not directly represented in the forest model, it is assumed that they can be obtained from the foliage (leaf) substrate concentrations, *C*_le_ and *N*_le_ [kg substrate C, substrate N (kg structural dry matter)^−1^] by simple multipliers:

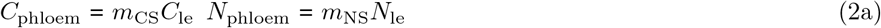

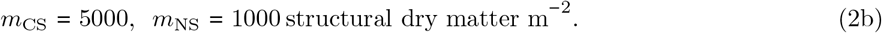

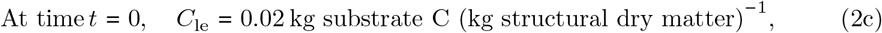

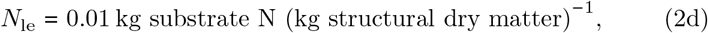

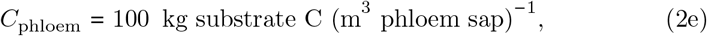

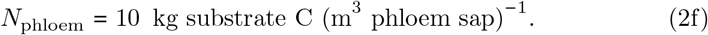

In Eq (2) the units of the phloem C and N concentrations, *C*_phloem_ and *N*_phloem_, are kg substrate C, substrate N (m^3^ phloem sap)^−1^ in the phloem. The values for the multipliers *m*_CS_ and *m*_NS_ are chosen so that reasonable values of phloem concentrations are obtained [5, 89]. For example, at time *t* = 0, *C*_le_ = 0.02 kg substrate C (kg structural dry matter)^−1^ converts to *C*_phloem_ = 100 (= 5000 × 0.02) kg substrate C (m^3^ phloem)^−1^ = 100 g substrate C (litre phloem)^−1^ = 100/144 mol sucrose litre^−1^ = 0.7 molar sucrose in phloem (assuming that sucrose is C_12_H_22_O_11_); *N*_le_ = 0.01 kg substrate N (kg structural dry matter)^−1^ converts to 10 (= 1000 × 0.01) kg N m^−3^ = 10 g N litre^−1^ = 10/14 = 0.7 molar glutamate (assuming that glutamic acid is C_5_H_9_NO_4_).

The phloem C:N substrate ratio [kg C (kg N)^−1^] is [with Eq (2)]

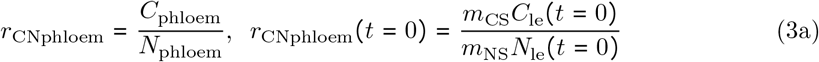

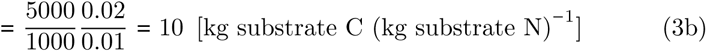

## Appendix D Aphid submodel

The aphid submodel is shown schematically in Fig 2. The model has 10 state variables. There are two morphs of the aphid: alate (winged; ala) and aperterous (wingless, apt). Each morph has four juvenile stages called instars, and an adult stage. All aphid state variables have units of numbers of aphids per stem. All notation is summarized in Appendix E.

We define the total number of alate instars (*a*_lai_), alate adults (*a*_laa_), the total alates (*a*_la_), the same for apterous aphids (*a*_pti_, *a*_pta_, *a*_pt_) and the total number of aphids (*a*_ph_) as:

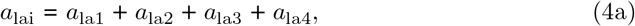

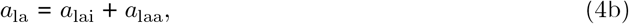

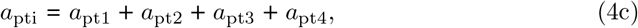

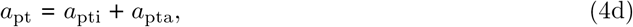

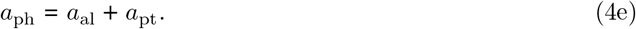

At *t* = 0, *a*_ph_ = 10 aphids per stem.

The spruce aphid does not colonize and feed on the current year’s foliage [90]. From the four foliage categories in the efm, only the last three are used here for the calculation of aphid densities on foliage (*A*_leaf,aph_). Dividing by the colonized leaf area per stem, *A*_leaf,aph_ (m^2^ stem^−1^), aphid density (ρ) per unit area of foliage (aphids m^−2^) are:

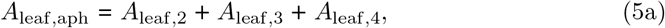

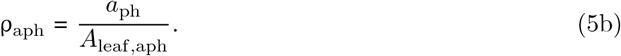

At *t* = 0, A_leaf,aph_ = 0.009 (m^2^ stem^−1^), and ρ = 10/0.009 = 1111.1 aphids (m^2^ leaf)^−1^. The *t* = 0 values are for young seedlings [Eq (1)] and the ‘standard’ aphid infestation, Eq (64) and for the plantation defined in Section 4.2 above. Note that an aphid density of 1000 aphids (m^2^ leaf)^−1^ corresponds to a distance between aphids of about 3 cm.

Using the aphid state variables, the C and N contents of the aphids per stem are (units: kg aphid C, N stem^−1^):

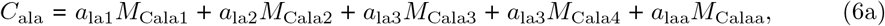

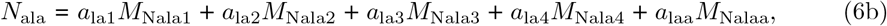

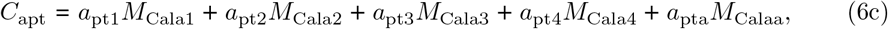

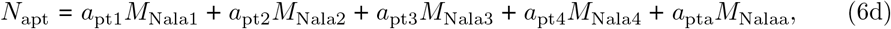

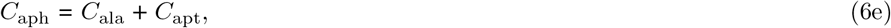

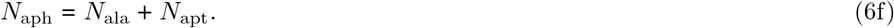

The C and N contents of each morphological aphid form (Fig 2) are constants, unaffected by growth temperature or nutritional status and are (units: kg C, N aphid^−1^):

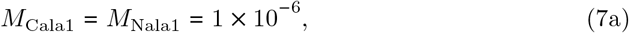

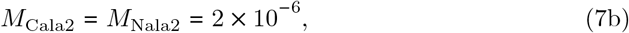

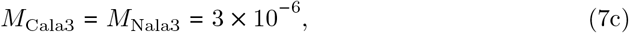

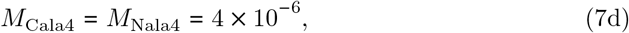

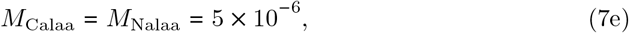

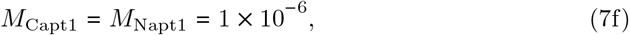

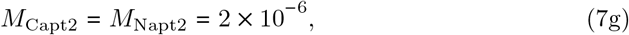

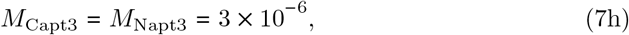

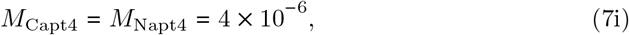

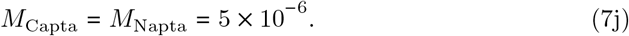

These estimates were made to enable the model to be completed, although we have been unable to find measurements of these quantities. The values are not critical for the range of behaviour exhibited by the model (See chapter 4 of Dixon [4] and particularly figure 4.1 on p. 59; note that 1 × 10 ^−6^ kg = 1 mg = 1000 μg).

### D.1 Apterous (wingless) adult aphids, *a*_pta_

#### D.1.1 Inputs: apterous (wingless) adult aphids

There is a single input, from the developmental output flux of the fourth apterous instar (*a*_pt4_, Fig 2), *O*_apt4→a_, calculated in Eq (42) (the output is assumed to become an input without loss). The input, *I*_apta_, requires C and N fluxes, of *I*_Capt4→a_ and *I*_Napt4→a_:

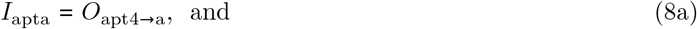

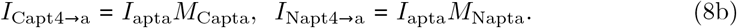

Units of the first of Eq (8) are aphids stem^−1^ d^−1^ and of the other two are kg aphid C, N stem^−1^ d^−1^. See Eq (7) for the C, N contents of apterous adults.

#### D.1.2 Outputs: apterous (wingless) adult aphids

It is assumed that there is no flux of aphids to surface litter accompanying foliage litter flux. Thinning, as applied in ‘plantation’ mode [Section 4.2, Eq (1)] — i.e. removal of whole trees (stems), does not affect aphid state variables which are aphids per stem (although aphid number per unit ground area is affected). This leaves two outputs: mortality (dependent on temperature and nutrition) and pruning (removing branches including foliage).

##### Mortality

Our approach to aphid mortality is mostly guess work, as there is little guidance on the topic which is relevant to a mechanistic modelling exercise where ordinary differential equations are the basic representational tool. Some authors prefer to talk of ‘survival’ (e.g. Duffy et al., 2017 [63]; see their figure 3), but survival then depends on a development rate (Section D.4) as well as a mortality rate. Dixon ([4], p. 165, see their figure 7.25) presents mortality rate as related to relative growth rate, without specifying units, rather than relating mortality rate to forest sub-model variables (such as phloem N) and driving variables (such as air temperature). We ignore any possible influence of predators on mortality. There is some evidence from other mechanistic aphid modelling that considering predation does not fundamentally effect the conclusions regarding the impacts of climatic change [29].

It is assumed that aphid mortality, k_aph,mort_, depends on air temperature, *T*_air_ and phloem N, *N*_phloem_; the specific rates (d^−1^) are calculated independently [Eqs (9) and (10)] and then combined in Eq (11). Adult apterous mortality rate is then *O*_apta→mort_ (aphids stem^−1^ d^−1^), given by Eq (12).

The specific air-temperature-dependent mortality rate, *k*_aph,Tmort_ (d^−1^), is given by a skewed inverted parabola (Fig D.1):

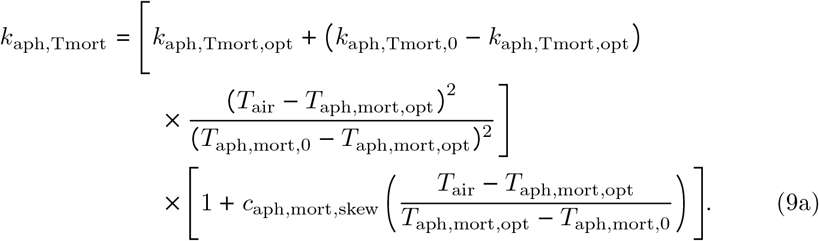

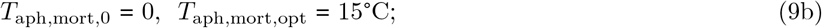

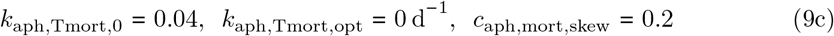

**Fig D.1.**
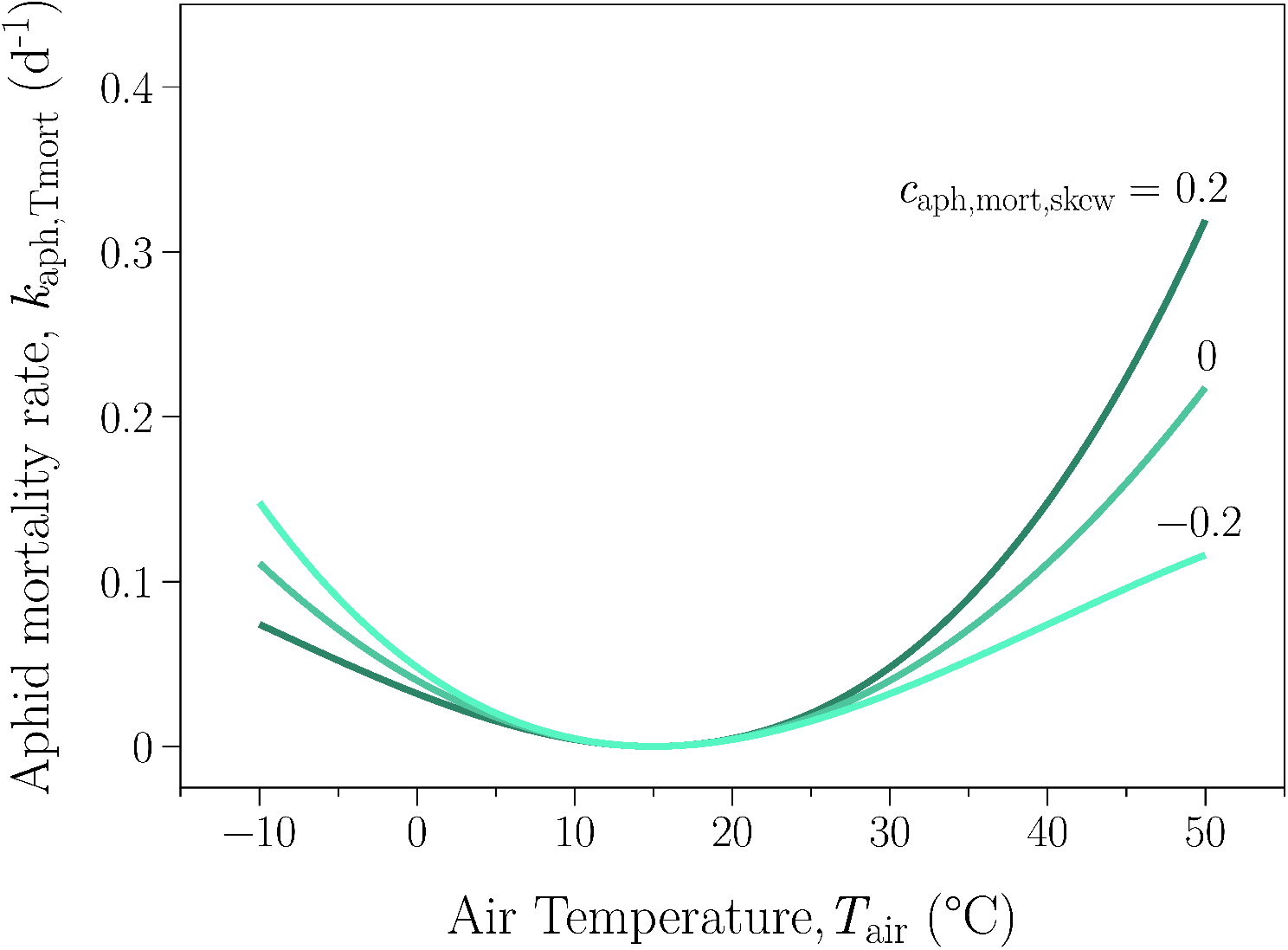
Aphid mortality rate as affected by temperature. Eq (9) is plotted for three values of the skewness parameter *c*_aph,mort,skew_ as given. The default value used in the simulations is 0.2, shown by the dark line. Other parameters are as in Eq (9).

If air temperature *T*_air_ equals the optimum temperature for minimum aphid mortality, *T*_aph,mort,opt_, then mortality rate *k*_aph,Tmort_ = *k*_aph,Tmort,opt_ is zero. If temperature *T*_air_ = *T*_aph,mort,0_ (the second reference point, taken here to be 0 °C), then *k*_aph,Tmort_ = *k*_aph,Tmort, 0_ = 4% d^−1^. The second term in square brackets skews the response about the optimum temperature (here 15 °C), according to the value of the skewness parameter *c*_aph,mort,skew_. With the value given, if *T*_air_ = *T*_aph,mort,0_, then the mortality rate is decreased by 20%. The temperature response of mortality rate is drawn in Fig D.1 for several values of the skewness parameter.

The most significant part of the response drawn in Fig D.1 is the increase in mortality as the temperature is lowered. This causes the aphid infestation to become less severe or even zero as the temperature is lowered (Fig 4).

Nutrition-dependent mortality rate, *k*_aph,Nmort_ (d^−1^), is described by a ‘switch-off’ sigmoidal response (e.g. [37], equation 4.61, figure 4.9, pp. 109-111), to phloem N concentration, *N*_phloem_ [Eq (2), Fig D.2], given by

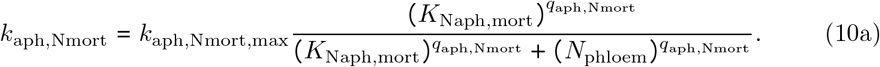

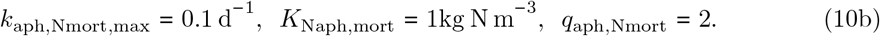

**Fig D.2.**
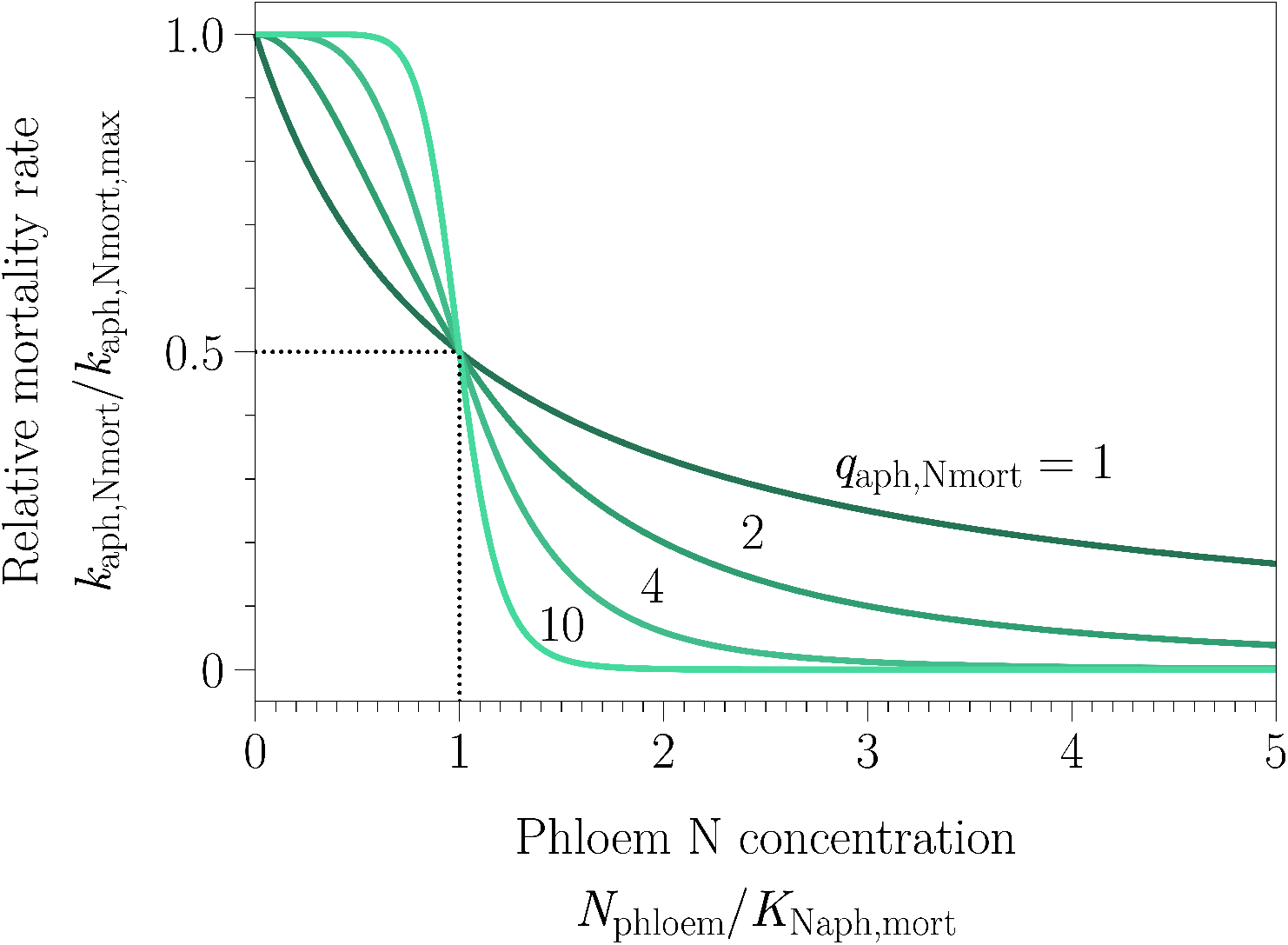
Nutritional response of aphid mortality. Aphid mortality rate as affected by N nutrition, in particular, N concentration in the phloem, *N*_phioem_ [Eq (2)]. Eq (10) is plotted for four values of the steepness parameter, *q*_aph,Nmort_, as given. The default value used in the simulations is *q*_aph,Nmort_ = 2.

The maximum mortality rate, *k*_aph,Nmort,max_, if there is no food (*N*_phloem_ = 0) is 10% per day. The half-maximal mortality rate (5% per day) occurs when *N*_phloem_ = *K*_Naph,mort_ = 1 kg N m^−3^ = 1/14 mol N. Note that, from Eq (2), this corresponds to a foliage substrate N concentration of *N*_le_ = 0.01 kg substrate N (kg structural dry matter)^−1^. The nutritionally-dependent mortality rate approaches zero at high values of *N*_phloem_. The steepness of the response depends on *q*_aph,Nmort_, here assigned the value 2. The response is drawn in Fig D.2 for four values of *q*_aph,Nmort_.

It is assumed that the overall mortality rate (d^−1^), k_aph,mort_, is given by summing the temperature-dependent and the nutritionally-dependent mortality rates [Eqs (9) and (10); assuming aphid temperature = air temperature, *T*_air_ (°C)]. Therefore:

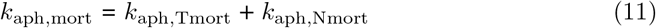

The mortality rate varies between 2% and 6% per day, depending on temperature and N nutrition (Fig D.3B).

**Fig D.3.**
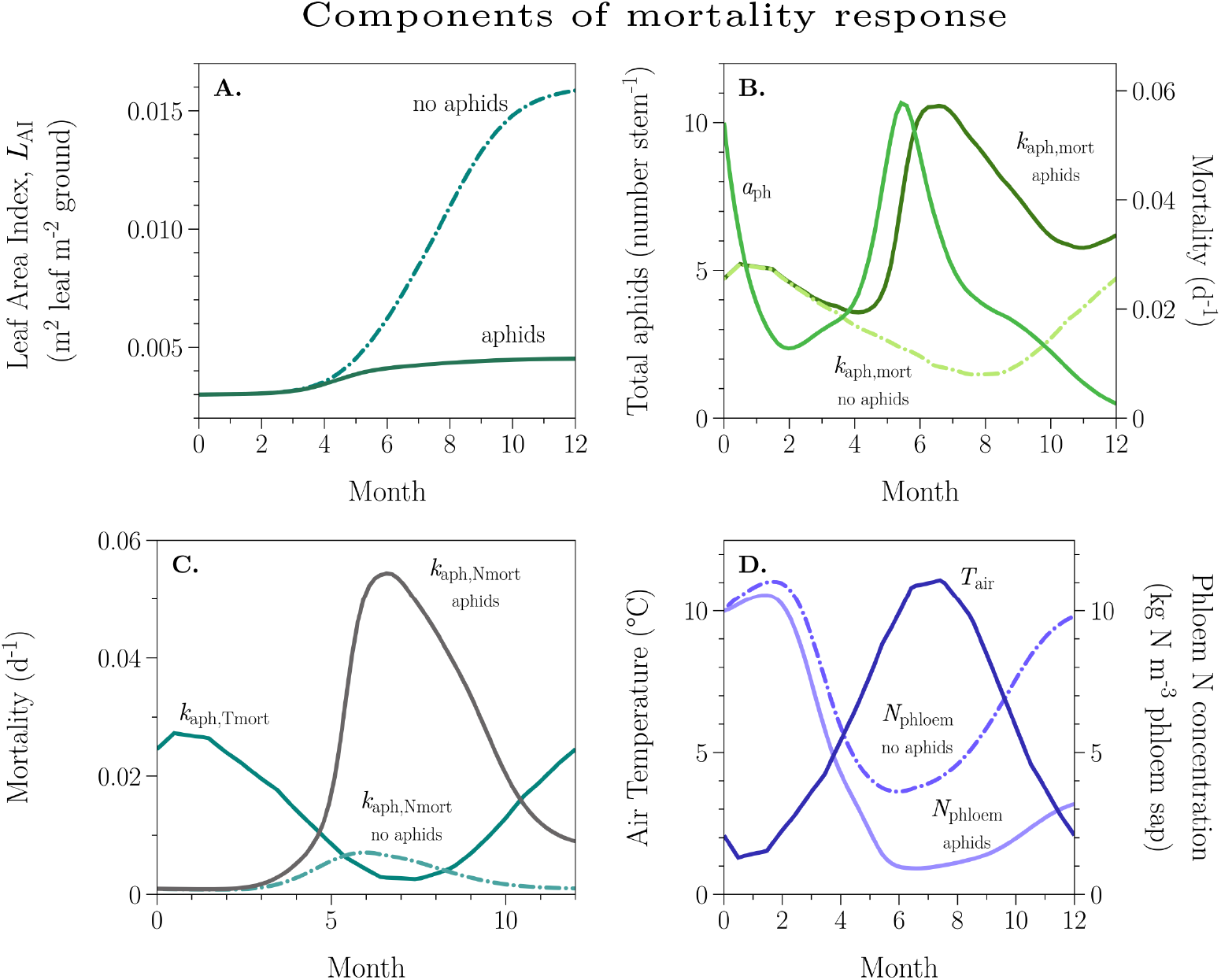
Aphid mortality. Contributions to aphid mortality are shown over 12 months without aphids being present (dash-dot lines) and with aphid infection at *t* = 0 (solid lines) of 10 adult alates per stem [Eq (64)]. At t = 0 d the seedling plants have an LAI of 0.003 and a stem density 0.25 stems m^-2^ [See also Eqs (1), (4) and (5)]. **A**, leaf area index, L^AI^ [Eq (1)]. **B**, total aphids, a_ph_ [Eq (4)]; aphid mortality rate, k_aph,mort_ [Eq (11)]. **C**, temperature-dependent mortality, k_aph_,T_mort_ [Eq (9)]; nitrogen-dependent mortality, kaph,Nmort [Eq(10)]. **D**, Air temperature, T_air_ (Fig 3), the determinant of temperature-dependent mortality in **C**; phloem N [Eq (2)], the determinant of nutrition-dependent mortality in **C**, phloem N is decreased by aphid infestation (solid line).

We have mentioned above (Section D.1.2), our difficulties with mortality rate. We use the same mortality rate, k_aph, mort_ [Eq (11)], for all the aphid state variables (Fig 2), although this could be easily relaxed. In Fig D.3, the components of mortality are examined. Mortality rate can be calculated in the absence of aphids, although the presence and activity of aphids depresses phloem N (Fig D.3D) and this increases N phloem-dependent mortality [*k*_aph,Nmort_, Fig D.3C, Fig D.2, Eq (10)].

Temperature-dependent mortality, k_aphTmort_, is the same whether aphids whether aphids are present or not [Fig D.3C, Eq (9)]. Fig D.3A shows the effect of aphids on leaf area index [Eq (1)] at a constant stem density, nstems, [Eq (1)]. Later, after considering developmental rates (Section D.4), we examine survival in Section D.4.1 and Fig D.8.

With Eq (11) for specific aphid mortality rate, *k*_aph,mort_ (*d*^−1^), output from the apterous adult pool, *a*_pta_, to mortality is (aphids stem^−1^ d^−1^)

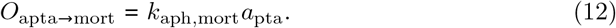

There are output (*O*) C and N fluxes associated with this aphid flux (using an obvious notation):

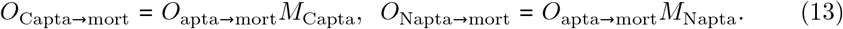

Units are kg C, N stem^−1^ d^−1^. Aphid C and N contents are given in Eq (7). These fluxes are input to the soil surface litter pools [Eqs (108) and (118)].

##### Pruning

The pruning flux of apterous adults is *O*_apta→prune_ (aphids stem^−1^ d^−1^). Foliage pruning, if applied, occurs at a rate of k_le→prune_ (d^−1^). This gives rise to outputs (losses) of aphids and associated fluxes of C and N (kg aphid C, N stem^−1^ d^−1^), of

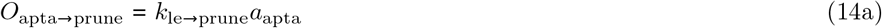

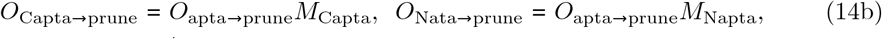

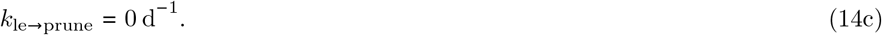

The foliage pruning constant, *k*_le→prune_, is not used in the simulations presented here and it is set to zero. This aphid output flux is added to the mortality flux in Eq (15).

The total output of adult apterous aphids, *O*_apta_, is obtained by adding the contributions from mortality [Eq (12)] and pruning [Eq (14)]:

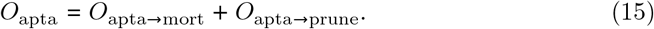

#### D.1.3 Differential equation: apterous (wingless) adult aphids

The differential equation (apterous aphids stem^−1^ d^−1^) and initial value for the state variable, *a*_pta_, for the apterous adult aphids is:

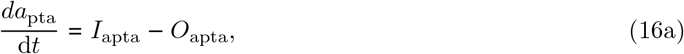

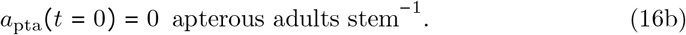

The input and output terms are given by Eqs (43) and (15).

### D.2 Apterous aphid fecundity

Adult aphids give birth to offspring or nymphs. The rate at which this occurs is influenced by temperature and phloem N concentration. The N ingested is entirely converted to wingless (apt) and winged (ala) 1st instar nymphs, *a*_pt1_ and *a*_la1_ (Fig 2). These can have different N contents [Eq (7)]. Account must be taken of this before total and fractional fecundities can be calculated [Section D.3.4; Eq (29)].

#### D.2.1 Apterous aphid fecundity as a function of temperature

A general and easily adjustable temperature response function, *f*(*T*), used for many plant and soil biological processes is ([37], pp. 105-106)

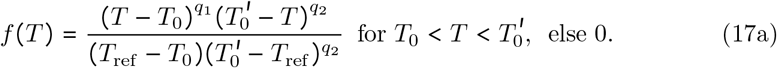

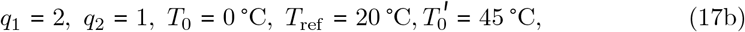

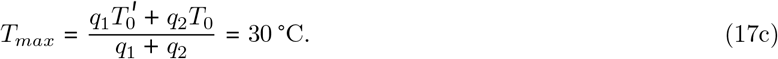

Here default values of the five parameters are given; the shape of the default response is illustrated by the continuous lines in Fig D.4A-D. The default is a cubic. *f*(*T*) is only non zero between temperatures *T*_0_ and 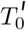. Fig D.4A shows the effect of varying the parameter *q*_1_. The response is initially sigmoidal if *q*_1_ > 1. In Fig D.4B, *q*_2_ is varied. It can be seen that the steepness of the high-temperature decrease is highly dependent on *q*_2_ for *q*_2_ < 1, although the early part of the curve is much less affected by *q*_2_. In Fig D.4C, *T*_0_ is varied and in Fig D.4D, 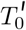. *T*_ref_ is a reference temperature: *f*(*T*) = 1 at *T* = *T*_ref_ = 20 °C. The five parameters define: the zero points of *f*(*T*) (*T*_0_ and 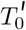); the shape of the curve in the neighbourhood of the zero points (*q*_1_ and *q*_2_); and the reference temperature, *T*_ref_. An equation for the temperature where *f*(*T*) is maximum is given, *T*_max_; the temperature of any point of inflexion can be derived.

**Fig D.4.**
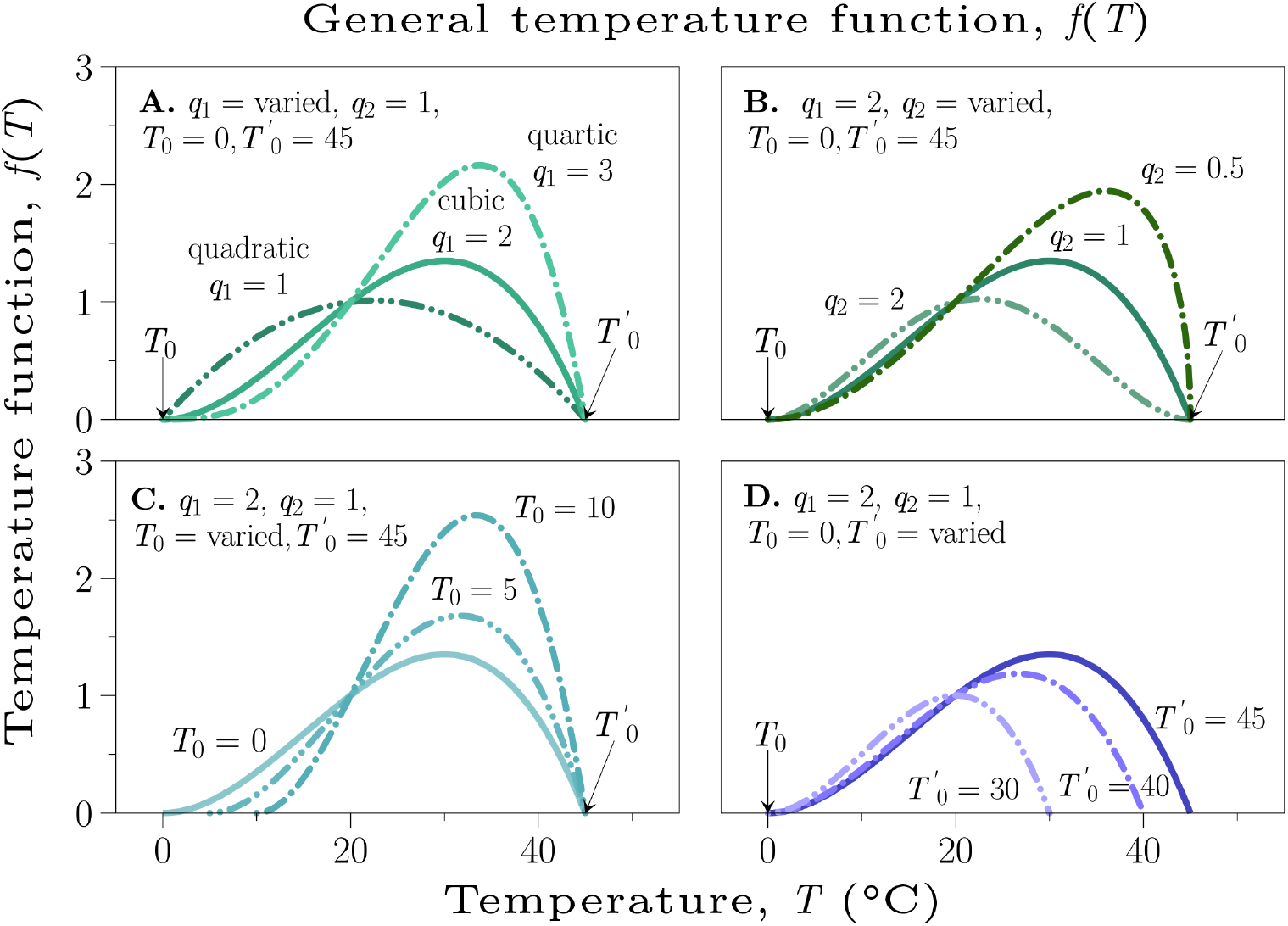
General biological temperature function. General biological temperature function, Eq (17), with five parameters: *q*_1_, *q*_2_, *T*_0_, 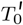 and *T*_ref_, of which four are varied in the figure. This is applied in the efm and the aphid sub-model for various processes. It is drawn, in A for three values of parameter *q*_1_, in B for three values of *q*_2_, in C for three values of *T*_0_ and in D for three values of 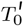. Otherwise, the parameters have the values given in Eq (17). Commonly used default values are shown by the continuous heavy lines.

Taking *T* = *T*_air_ and with the default parameters given, Eq (17) is used to modify many above-ground (shoot) plant processes (where it is reasonable to take shoot temperature equal to air temperature, *T*_air_), as well as the temperature dependence assumed for some aphid processes, e.g. emigration [Eq (61)].

The effect of temperature on aphid fecundity, *f*_Taph,fec_, is obtained by using air temperature, *T*_air_ and the standard temperature function in Eq (17) and Fig D.4, but with two different temperature parameters (*T*_0_, 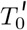):

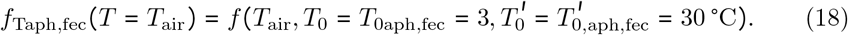

This is drawn in Fig D.5A. It can be compared with Duffy et al.’s [63] figure 2, who make use of data from Dean [91], although our calculations of total and fractional fecundities are completed below [Section D.3.4, Eq (29)].

**Fig D.5.**
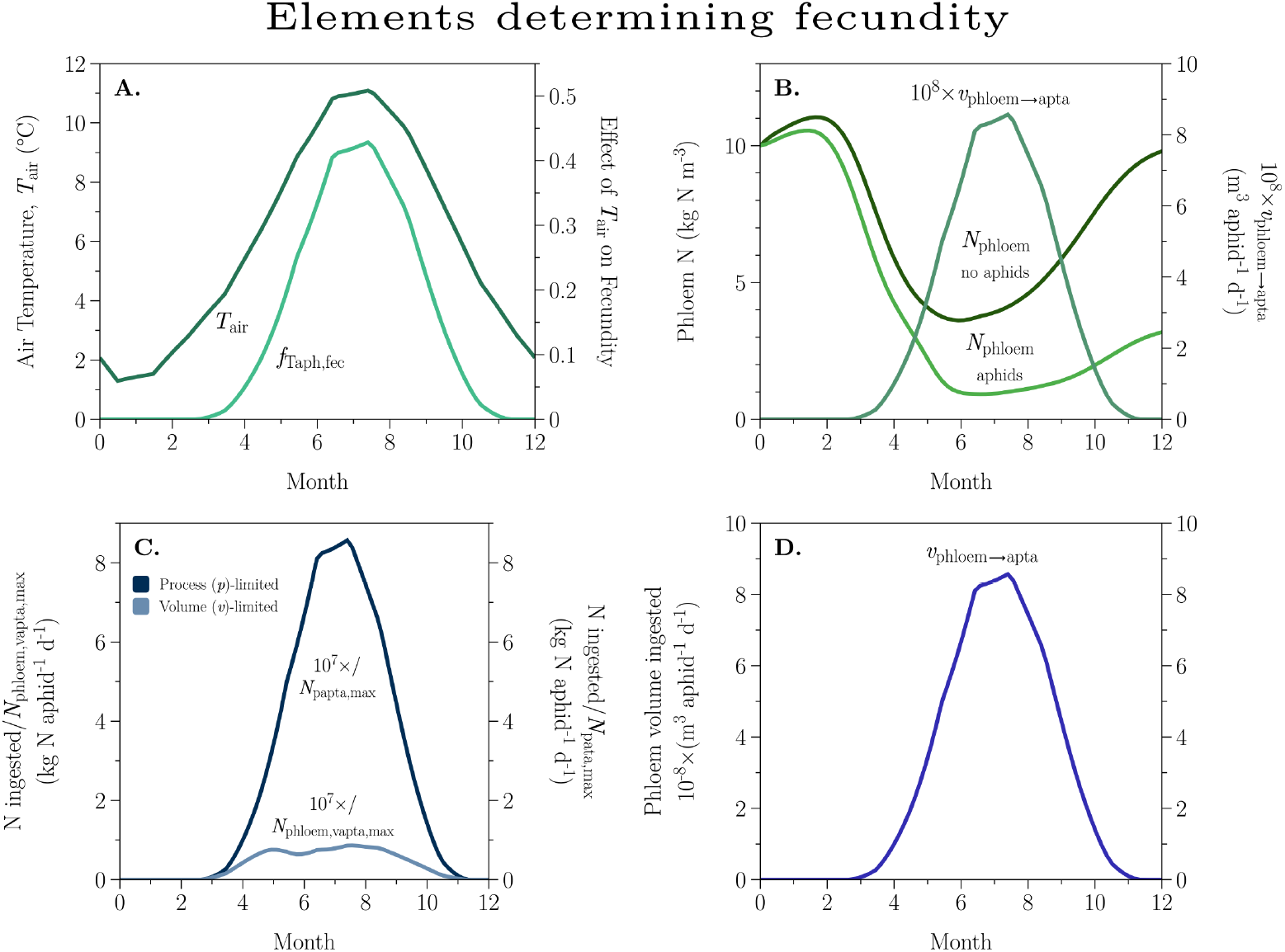
Elements determining fecundity. Factors contributing to fecundity are shown as they occur in an Eskdalemuir environment (**Section 4.1**), for the first year of growth of typical spruce plantation (**Section 4.2**) infected with ten alate adults at time zero [**Eq (64)**]. A, air temperature, T_a_ļ_r_, and its effect on fecundity, *f*_Taph,fec_ [**Eq (18)**]. B, phloem N concentration, *N*_phioem_ [**Eq (2)**] is illustrated without and with aphid infection which lowers *N*_phioem_ levels and thereby increases the actual volume of phloem sap ingested, *v*_phioem→apta_ [**Eq (23)**]. C, alternatives for the N ingested per aphid per day: (i) process-limited (*p*) N intake, *I*_Npapta,max_ [**Eq (21)**]; (ii) phloem-volume-limited (*v*) N intake, *I*_N_phloem,vapta,max__ [**Eq (20)**]. D, actual phloem volume ingested per aphid per day, *v*_phloem→apta_ [**Eq (23)**].

#### D.2.2 Apterous aphid fecundity as a function of phloem N concentration

Aphids are assumed to have a maximum volume intake of phloem sap at a given temperature, *v*_apta,max_ (m^3^ aphid^−1^ d^−1^), obtained by f_Taph, fec_ (*T*_air_) of (Fig D.5A) multiplied by the maximum volume intake at the reference temperature of 20 °C:

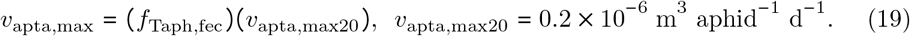

The 20 °C value, *v*_apta,max20_, corresponds to 0.2 ml aphid^−1^ d^−1^. The maximum phloem-volume limited N intake per aphid is (units are kg N aphid^−1^ d^−1^) (Fig D.5C):

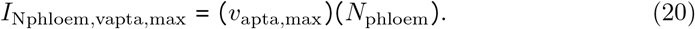

*N*_phloem_ is given by Eq (2) [kg N substrate in phloem (m^3^ phloem sap)^−1^]. This may exceed the N requirement of the aphid, defined as the maximum amount of N which can be processed (*p*) at the ambient temperature (*T*_air_), calculated by

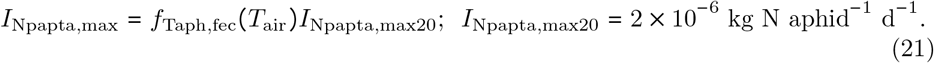

*f*_Taph,fec_(*T*_air_) is given in Eq (18). *I*_Npapta,max20_ is the maximum amount of N that can be processed (*p*) at the reference temperature of 20 °C. *I*_Npapta,max_ is illustrated in Fig D.5C. Note that here we have used the same temperature dependence, *f*_Taph,fec_, for the process-dependence [Eq (21)] as for the volume limitation [Eq (19)].

The actual amount of N ingested per aphid per d is the least of the volume-intake-limited N-intake [Eq (20)] or the processing-limited N-intake [Eq (21)] (kg N aphid^−1^ d^−1^):

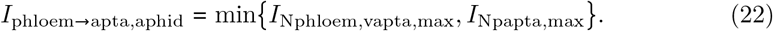

Neither of the two arguments is affected by the presence of aphids—e.g. which will depress *N*_phloem_ (Fig D.5B). In fact the processing-limited maximum is about eight-times as large as the volume-limited maximum (Fig D.5C), so the volume-limitation wins the argument in Eq (22).

Dividing by *N*_phloem_ [Eq (2)], Fig D.5B] (kg N m^−3^), which is affected by the aphid infestation, the actual volume of phloem sap ingested per apterous adult aphid is (m^3^ aphid^−1^ d^−1^):

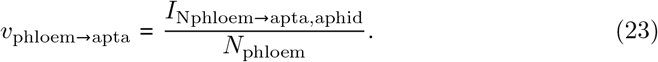

This is illustrated in Fig D.5B and Fig D.5D.

The C intake per aphid (kg C aphid^−1^ d^−1^) and the N and C intakes per stem (kg N, C stem^−1^ d^−1^) are respectively:

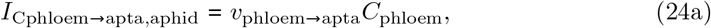

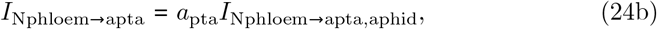

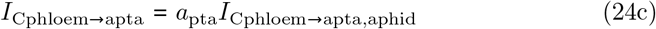

The right side quantities are given in Eqs (23), (2), (16), (22) and the first of (24).

### D.3 Alate:apterous ratio in offspring from apterous adults

The N ingested is assumed to be completely converted to wingless (apt) and winged (ala) first instar nymphs, in pools *a*_pt1_ and *a*_la1_ (Fig 2). Because these two forms can have different masses (N contents) [Eq (7)], the fractions of each type must first be calculated before total fecundity can be determined (Section D.3.4). It is assumed that the ratio is affected by three factors: temperature, total aphid density and nutritional status [92]. It is assumed that the three factors operate multiplicatively and for each factor x, the fraction of the offspring of apterous female adults which are alate (winged), *f*_x,apt→ala_, is calculated. The possible effects of day length are ignored (but see figure 6.8, p. 109 in [4]). Duffy et al. ([63], equation 4) assume the percentage of nymphs which are alates increases with aphid density and also growth stage of the cereal crop.

#### D.3.1 Alate:apterous ratio as a function of temperature

Higher air temperatures, *T*_air_ (°C), generally give more alates ([4], page 109). A positive (switch-on) sigmoidal dependence on air temperature (*T*_air_) is assumed ([37], pp. 109 - 110), with:

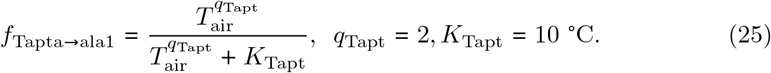

This is drawn in Fig D.6. *K*_Tapt_ is the half-maximal response temperature. *q*_Tapt_ determines the steepness of the response; *q*_Tapt_ = 1 gives the familiar Michaelis-Menten response.

#### D.3.2 Alate:apterous ratio as a function of aphid total density

This is ρ_aph_ [aphids (m^2^ leaf area)^−1^], Eq (5). Higher aphid densities cause more of the apterous offspring to be alates (winged). As in Eq (25) and Fig D.6 above for relative air temperature, a positive sigmoidal dependence on aphid density, ρ_aph_, is assumed:

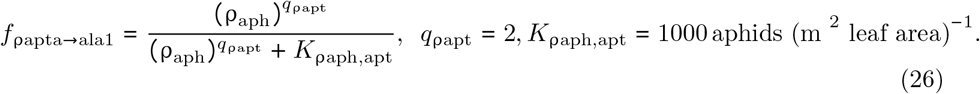

**Fig D.6.**
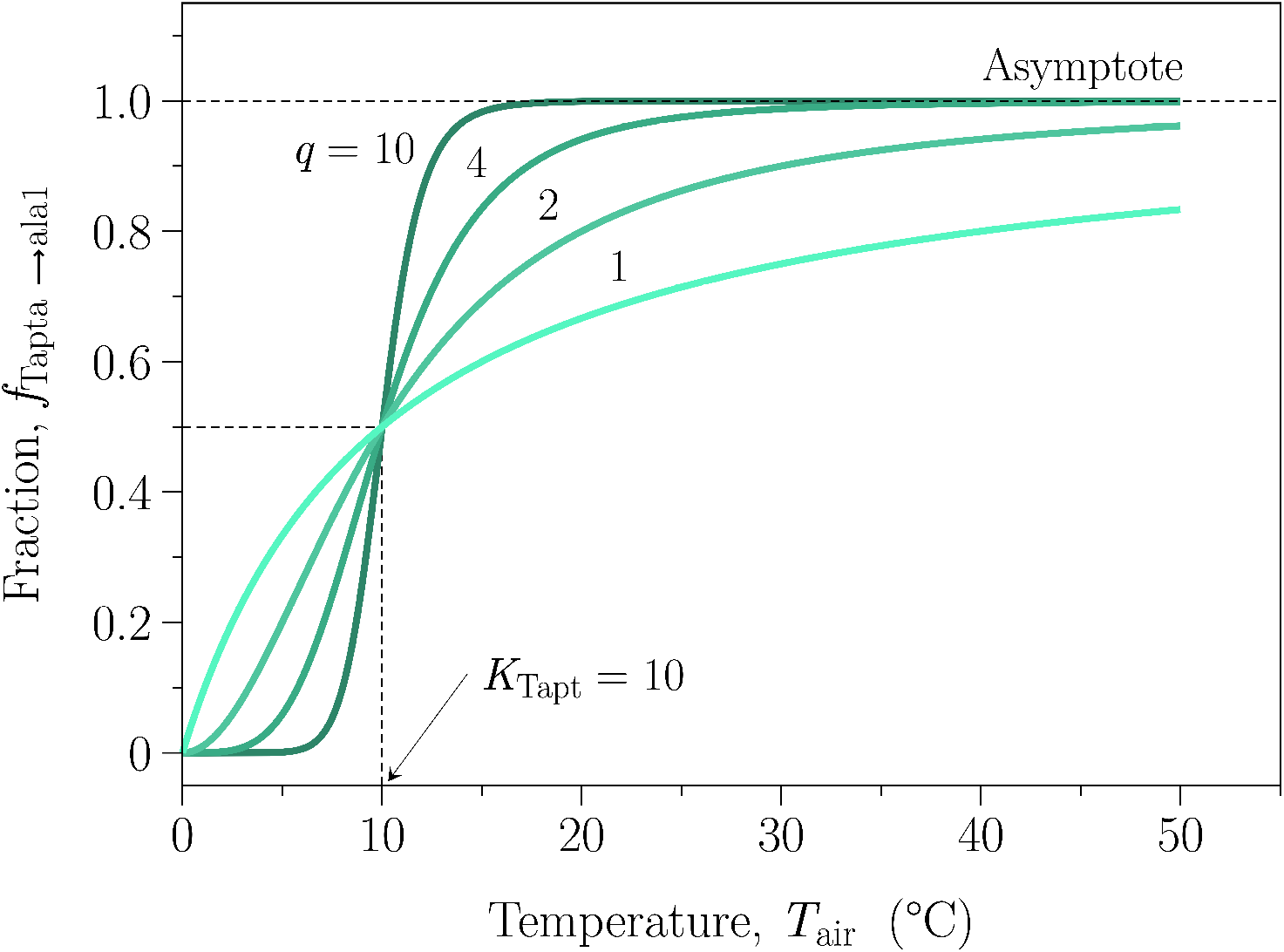
Alate fraction of apterous offspring as affected by temperature. Eq (25) is drawn for four values of the *q* parameter. The default value used in the simulations is *q* = 2.

The half-maximum density point is where ρ_aph_ = *K*_paph,apt_; the value given is equivalent to aphids being on a square grid 3 cm apart and is of the same order as the annual peak in ρ_aph_.

#### D.3.3 Alate:apterous ratio as a function of nutritional status

In this case, good nutritional status [a high value of *N*_phloem_, Eq (2)] gives few alates and the alate fraction approaches zero for high values of *N*_phloem_. Low nutritional status and a low *N*_phloem_ causes the alate fraction *f*_Napta→ala1_ to approach unity. An expression similar to that used in Fig D.2 for aphid mortality is employed [a ‘switch-off’ sigmoid rather than the ‘switch-on’ sigmoid used in Eq (26) and Eq (25); Fig D.6] ([37], figure 4.9, p. 111):

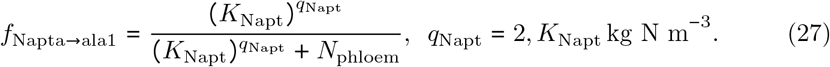

The half-maximal phloem N concentration of *K*_Napt_ = 4 kg N m^−3^ is equivalent to a foliage N substrate concentration of *N*_le_ = 0.04 kg N substrate (kg structural dry matter)^−1^ [Eq (2)].

Combining these three factors multiplicatively [Eqs (25), (26) and (27)], therefore the fraction of offspring from apterous adults which are alate (winged) and are apterous are given by:

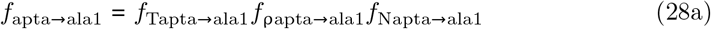

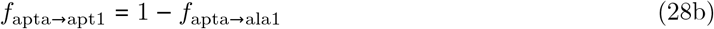

These three components and the outcome for *f*_apta→ala1_ are illustrated in Fig D.7.

**Fig D.7.**
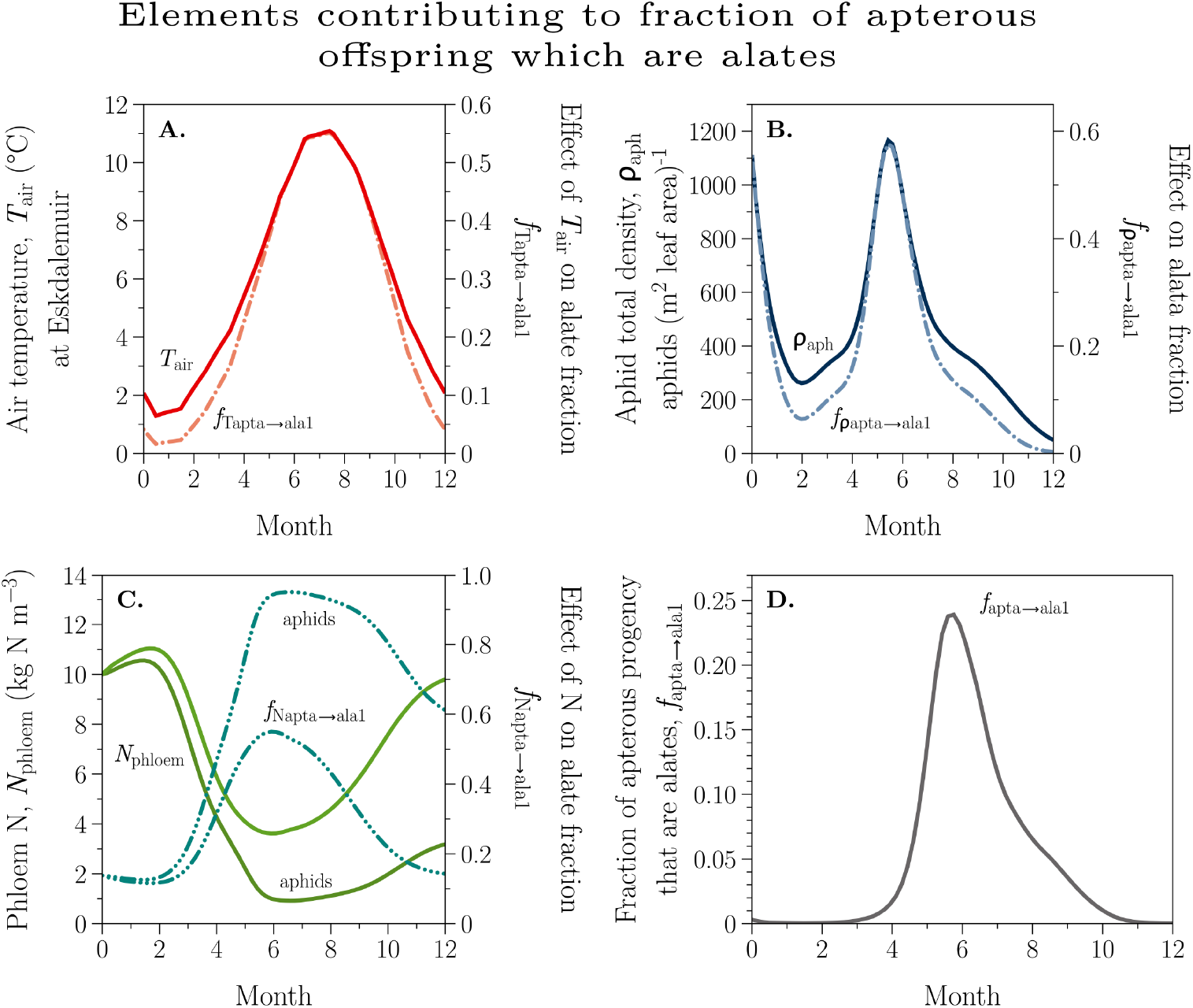
Elements contributing to the fraction of apterous offspring which are alates. Illustration of how the three different components contribute to the fraction of apterous offspring which are alates, as in Eq (28). This is for an Eskdalemuir environment (Section 4.1) and the first year of growth of typical spruce plantation (Section 4.2) infected with ten alate adults at time zero [Eq (64)]. **A**, air temperature, *T*_air_, via Eq (25). **B**, aphid density, ρ_aph_, via Eq (26). **C**, phloem N, N_phioem_, via Eq (27). Here, the introduction of aphids at time zero, depresses *N*_phioem_, the lower continuous line, resulting in a higher fraction destined for alates (upper dashed line). **D**, the three factors shown in **A**, **B**, **C** are combined in Eq (28).

The relative contributions to the fraction of apterous offspring assigned to the alate pathway are shown in Fig D.7 [Fig 2, Eq (28)]. In Fig D.7A, with dependence on air temperature, *T*_air_, this fraction, *f*_Tapta→ala1_ [Eq (25)] is not affected by the presence of aphids. In Fig D.7B, showing dependence on aphid density, ρ_aph_, the fraction is zero in the absence of aphids [Eq (26), with *ρ*_aph_ = 0]. The dependence on phloem N, *N*_phloem_, is illustrated in Fig D.7C [Eq (27)]. Because phloem N is consumed by aphids, their presence lowers *N*_phloem_, giving the lower continuous line for *N*_phloem_; this also raises the alate fraction of apterous offspring (upper dashed line). Last, in Fig D.7D, the three fractions shown in Fig D.7A–C are multiplied in Eq (28) to give the final allocation fraction, *f*_apta→ala1_.

#### D.3.4 Total fecundity and the fractional fecundity of apterous adults

Now that we have calculated the fractions of offspring from apterous adults which are alate and which are apterous, *f*_apta→ala1_ and *f*_apta→apt1_ in Eq (28), we are able to calculate the total fecundity of apterous adults. It is assumed that N ingested by apterous adult females per stem per day, *I*_N_phioem→apta__ [Eq (24)], is converted without loss into first-instar nymphs, giving a total rate of output to first instar offspring of *O*_apta→ala1_ + *O*_apta→apt1_, denoted by *O*_apta→axx1_ [with Eq (28)], of (aphids stem^−1^ d^−1^)

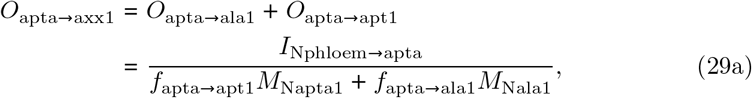

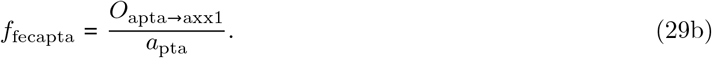

The aphid N contents are given in Eq (7). Equation (29b) defines the total fractional fecundity of apterous adults, *f*_fecapta_ (d^−1^); this is the total output of nymphs from apterous adults (*a*_pta_) towards the alate and apterous first-instar pools divided by the number of apterous adults (*a*_pta_). The transfer of nymph outputs into inputs into the first-instar pools is assumed to occur without loss giving (aphids stem^−1^ d^−1^)

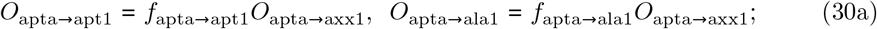

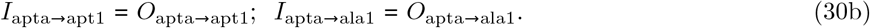

#### D.3.5 Associated N and C fluxes

There are output N fluxes from the *a*_pta_ pool associated with these aphid fluxes of (kg N stem^−1^ d^−1^):

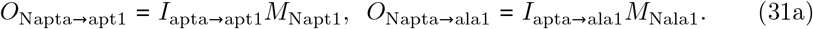

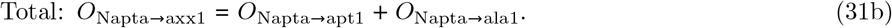

The right-side quantities for the first two equations are in Eq (30) and Eq (7).

There are carbon fluxes from apterous adults to respiration, offspring and honeydew [input to the surface litter metabolic pool, Eq (107)]. The carbon intake of the apterous adults is *I*_Cphloem→apta_ [Eq (24), kg C stem^−1^ d^−1^]. It is assumed that a fraction, *f*_Caph,resp_, of this is respired, giving a respiratory flux from apterous adults of (kg C stem^−1^ d^−1^)

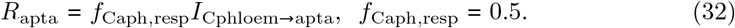

This value of 0.5 is typical of respiration losses [87].

Some carbon is used for first instar offspring (kg C stem^−1^ d^−1^):

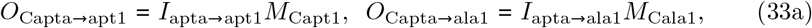

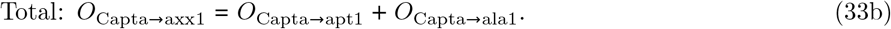

Here we have used Eqs (7) and (30).

The rest is honeydew (kg C stem^−1^ d^−1^) [with Eqs (24), (32) and (33)]

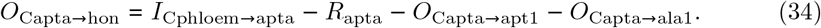

This output is input to the surface litter metabolic pool [Eq (107)].

### D.4 Development rates of apterous aphids

These depend principally on temperature (*T*_air_) and to a lesser extent on nutrition [*N*_phloem_, Eq (2)]. Aphid density does not seem to be a factor. We assume that day length is not important. For the temperature modifier of aphid development, *f*_Taph,dev_, the standard biological temperature response function is assumed as in Eq (17) but with one different parameter, *T*_0_ (Fig D.4D):

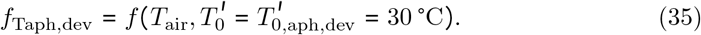

The Fig D.4D 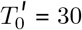 curve can be compared qualitatively with Duffy et al. ([63], their figure 3; see Section D.4.1 below).

The nutritional effect is small: a 10% decrease in development rate if *N*_phloem_ [Eq (2)] is zero so that *f*_Naph,dev,min_ = 0.9. The nutritional modifier of developmental rate, *f*_Naph,dev_, is assumed to be given by a Michaelis-Menten term with:

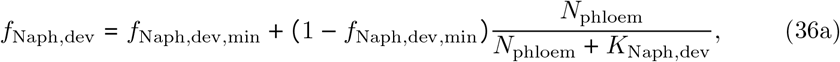

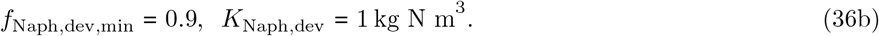

The *K* value, giving half-maximal nutritional effect, is equivalent to foliage N substrate concentration of *N*_le_ = 0.01 kg N substrate (kg structural dry matter)^−1^ [Eq (2)]. If *N*_phloem_ = 0, *f*_Naph,dev_ = 0.9; if *N*_phloem_ = *K*_Naph,dev_, *f*_Naph,dev_ = 0.95; *N*_phloem_ is large, *f*_Naph,dev_ = 1. Note that in the Fig 4 +0 °C run, *f*_Naph,dev_ is mostly about 0.97 + 0.02 and *N*_phloem_ is in the range 2 to 7 [Eq (2)] so the effect of nutrition on developmental rate is small.

The combined effects of temperature [Eq (35), Fig D.4D, 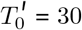 °C curve] and N nutrition on development [Eq (36)] are

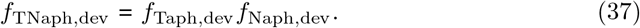

All the 20 °C apterous rate constants (d^−1^) are modified by this dimension-free factor:

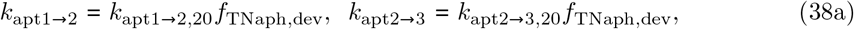

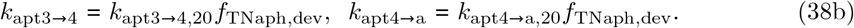

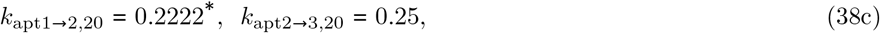

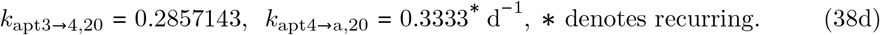

The lifetimes, *τ*(d), associated with these 20 °C rate parameters are

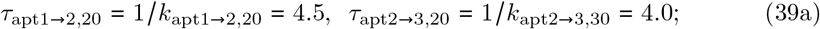

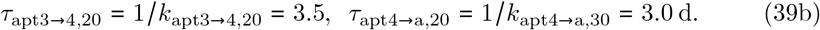

Summing, over all *τ*_apt1→a,20_ = 15 d.

The overall value for the transition from 1st apterous instar to apterous adult agrees reasonably with Dixon ([4], p. 109, table 6.1). Our attempts to relate present assumptions to the data, equations and figures of Duffy et al. [63] were unsuccessful. It is assumed that alates develop 15% more slowly than apterous aphids [Section D.9, Eqs (9) and (80)].

#### D.4.1 Survival

As mentioned in the first paragraph of Section D.1.2, survival is a concept used by some authors (e.g. [63]). In this subsection we show how it is related to the present formulation.

Considering the first instar apterous aphids, *a*_apt1_, the rate of transfer (units d^−1^) to the 2nd instar compartment is *k*_apt1→2_ [Eq (38), Fig 2]. This competes with aphid mortality, *k*_aph,mort_ [Eq (11), Fig D.3A], resulting in a survival probability which is a function of time, survival (*t*), whose asymptote is survival (*t* → ∞). Both quantities are dimensionless.

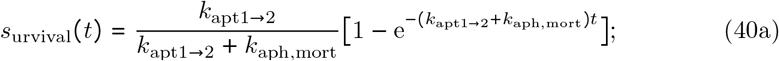

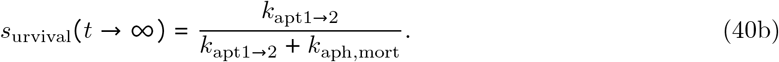

The latter is drawn in Fig D.8B and the quantities determining this in Fig D.8A.

**Fig D.8.**
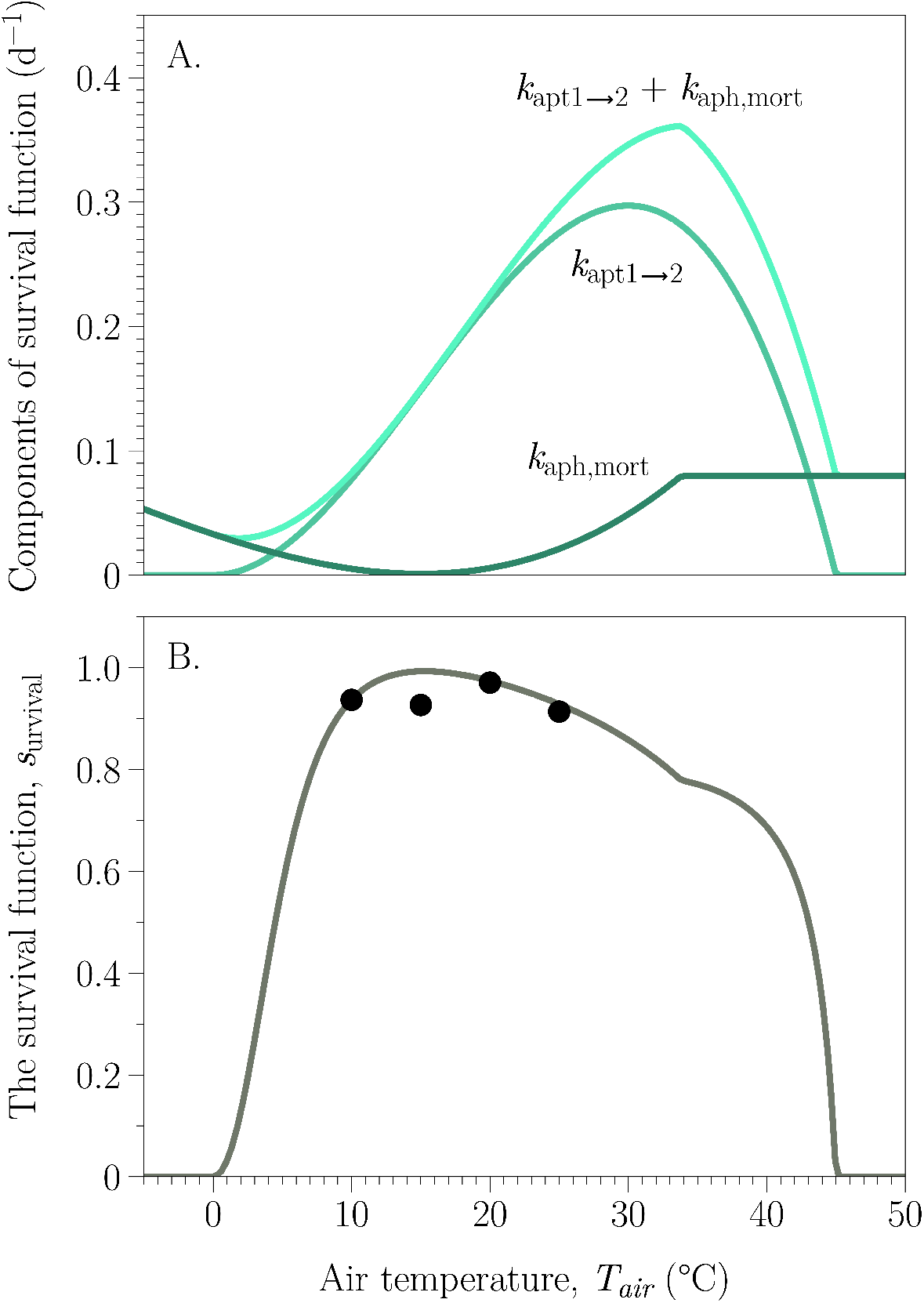
Survival. The temperature dependence of the survival function, Eq (40). **A**, the terms making up the function are shown: *k*_apt1→2_ is the specific rate of transfer of aphids from the 1st instar apterous compartment to the second [Fig 2, Eq (38)]; *k*_aph,mort_ [Eq (11), **A**] is the specific aphid mortality rate. **B**, the asymptotic survival function *s_urvival_*(*t* → ∞) [Eq (40)]. The four points shown are from Duffy et al. ([63], their figure 3), whose data are from Dean [91].

Figure 3 of Duffy et al. [63] compares reasonably with the present formalism and Fig D.8B. The survival fraction is a consequence of two competing rate constants, mortality and development. It may therefore be more pertinent to consider the underlying rate constants shown in Fig D.8A.

### D.5 Juvenile apterous (wingless) aphids

It is convenient to deal with these pools together (Fig 2). The four instars have state variables, *a*_pt1,…, 4_ (aphids per stem). First the pool inputs are defined, then the outputs and last the differential equations.

#### D.5.1 Inputs: juvenile apterous (wingless) aphids

These are: *I*_apt*j*_, *j* = 1,…, 4 (aphids stem^−1^ d^−2^), with (Fig 2)

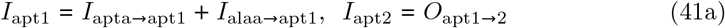

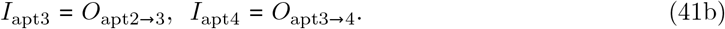

The inputs to the first pool are from the reproduction of both apterous and alate adults [Fig 2; Eqs (30) and (74)]. The inputs to the other three pools are equal to the developmental outputs of the preceding pools [Eq (42)], assuming that transfer takes place without loss.

#### D.5.2 Outputs: juvenile apterous (wingless) aphids

The outputs, *O*_apt*j*_, *j* = 1,… 4 (aphids stem^−1^ d^−1^) from each pool arise from development, mortality, tree thinning and/or foliage pruning. There is no loss of aphid numbers per stem to thinning when whole stems are removed.

##### Development

Developmental outputs are (aphids stem^−1^ d^−1^)

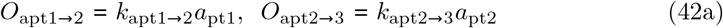

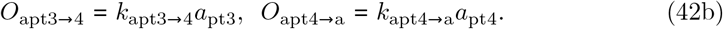

The developmental rate constants (d^−1^) are given in Eq (38). The state variables, *a*_pt*j*_, *j* = 1,…, 4 (aphids stem^−1^) are defined by Eq (47) (Fig 2). The outputs are transferred without loss, providing the inputs (aphids stem^−1^ d^−1^) of *I*_apt2-4_ in Eq (41) and also the input to apterous adults of

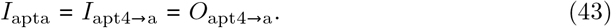

##### Mortality

It is assumed that the same mortality rate constant, *k*_aph,mort_ [Eq (11)], applies to all four instars, leading to outputs of (aphids stem^−1^ d^−1^)

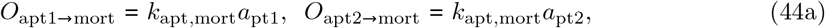

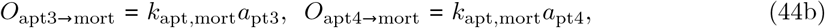

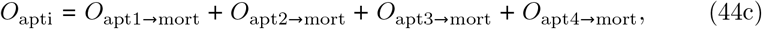

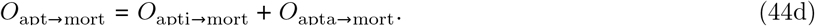

In the last two lines we calculate total instar mortality, *O*_apti→mort_ and total apterous aphid mortality, *O*_apt→mort_.

##### Pruning

Pruning is assumed to remove aphids from the aphid number per stem pools, as in Eq (14). The output fluxes are (aphids stem^−1^ d^−1^)

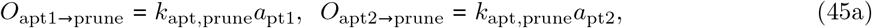

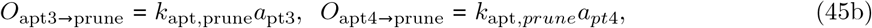

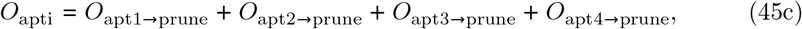

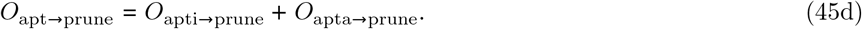

The state variables, *a*_pt*j*_, *j* = 1,…, 4 (aphids stem^−1^) are defined by Eq (47) (Fig 2). The last two of Eq (45) give the total apterous instar output to pruning and with the first of Eq (14), total outputs of apterous instars and total apterous aphids to pruning. See Section D.1.2 ‘mortality’. The pruning rate constant *k*_le→prune_ = 0 d^−1^ in these aphid studies [Eq (14)].

Combining Eqs (42), (44) and (45), the total outputs from the four pools are (aphids stem^−1^ d^−1^):

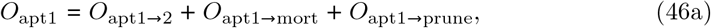

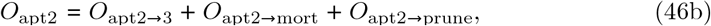

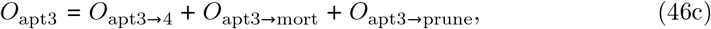

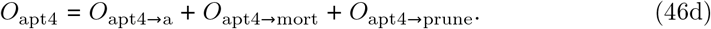

#### D.5.3 Differential equations: juvenile apterous (wingless) aphids

With Eqs (41) and (46), the differential equations (aphids stem^−1^ d^−1^) and initial values for the four apterous instar state variables are (aphids stem^−1^):

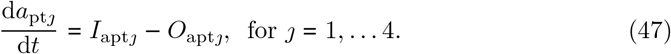

At *t* = 0 d: *a*_pt*j*_ = 0 for *j* = 1,…, 4 aphids stem^−1^.

#### D.5.4 Nitrogen and carbon fluxes for the four apterous instar pools

It is assumed that the N taken up from the phloem is the amount required to increase the N component of mass of the aphids which enter the next pool. Therefore, using an obvious notation (kg N stem^−1^ d^−1^):

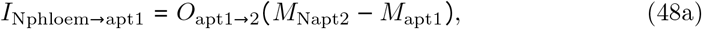

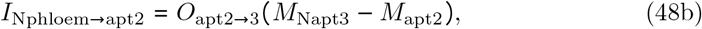

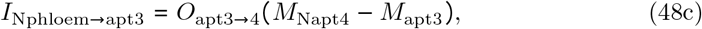

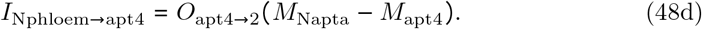

Aphid N contents are given in Eq (7); developmental outputs in Eq (42). Total N intake from the phloem by the four apterous instars (apti) is (kg N stem^−1^ d^−1^):

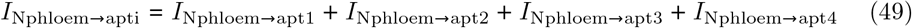

The C flux accompanying this N flux is (kg C stem^−1^ d^−1^)

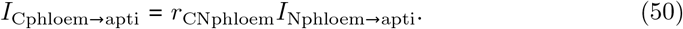

The phloem C:N ratio is given in Eq (3).

The C taken from the phloem is used for respiration, increase in instar mass and honeydew. The earlier treatment is followed [Eqs (32)–(34)].

The amount respired is (kg C stem^−1^ d^−1^)

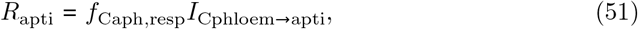

where fraction, *f*_Caph,resp_, is defined by Eq (32).

The increase in instar mass from aphid development (dev) also requires C from the phloem. The four input (*I*) components of this and the total phloem C output to supply these four inputs are (kg C stem^−1^ d^−1^)

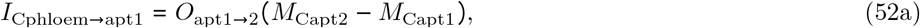

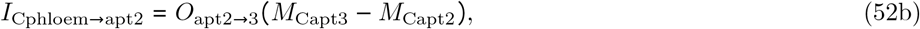

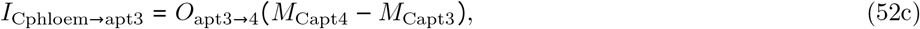

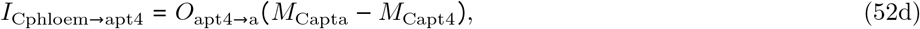

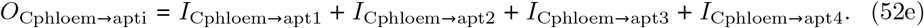

Equations (42) and (7) are used in the above.

Excess C is excreted as honeydew (hon), which is input to the surface litter metabolic pool [Eq (107)] ([34], chapter 5; kg C stem^−1^ d^−1^):

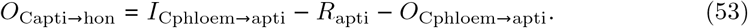

Eqs (50), (51) and (52) are used here.

Mortality (mort) gives rise to C and N outputs of (kg C, N stem^−1^ d^−1^):

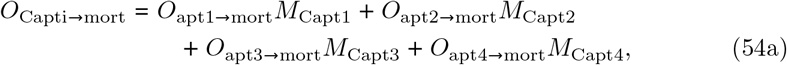

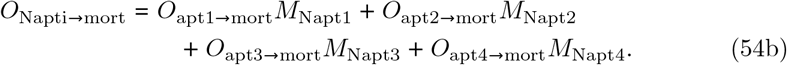

Equations (44) and (7) have been applied. These fluxes are input to the soil surface litter pools [Eqs (108) and (118)].

Pruning similarly gives rise to C and N outputs of (kg C, N stem^−1^ d^−1^):

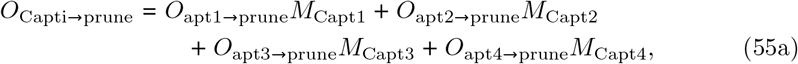

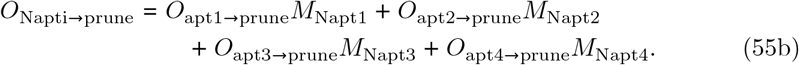

Equations (45) and (7) have been used, but see Eq (14).

### D.6 Alate (winged) adult aphids, *a*_laa_

#### D.6.1 Inputs: alate (winged) adult aphids

In this case [*cf*. Eq (8) and above], there are potentially two inputs. The first is from development of the 4th alate instar, *a*_la4_; this input, *I*_ala4→alaa_, is calculated in Eq (83). The second is from immigration, currently equated to zero [but see Eq (64) where *a*_laa_ is set to a non-zero value]:

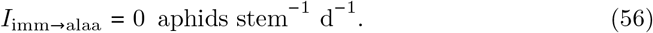

The total input to the alate adult pool is (aphids stem^−1^ d^−1^)

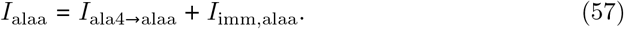

This requires C and N fluxes (kg C, N stem^−1^ d^−1^) of [*cf*. Eq (8)]:

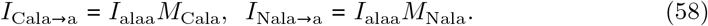

We have used Eq (7).

#### D.6.2 Outputs: alate (winged) adult aphids

There is assumed to be no aphid loss accompanying the foliage litter flux. Thinning does not affect aphid numbers per stem. There are outputs to mortality, pruning and emigration.

##### Mortality

The aphid flux to mortality, *O*_alaa→mort_ (aphid number stem^−1^ d^−1^), is assumed to depend on temperature and nutrition. The specific rate constant, *k*_aph,mort_ (d^−1^) assumed to be the same for all aphid morphs, is calculated in Eq (11). The output from the adult alate pool to mortality and the accompanying C and N fluxes are [*cf*. Eqs (12) and (13)]

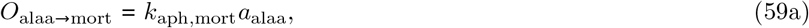

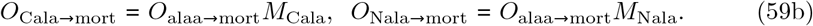

Aphid C and N contents are in Eq (7). Units are aphids stem^−1^ d^−1^ and kg C, N stem^−1^ d^−1^. These fluxes are input to the soil surface litter pools [Eqs (108) and (118)].

##### Pruning

The pruning flux, *O*_alaa→prune_ (aphids stem^−1^ d^−1^) and the associated fluxes of C and N, are, after Eq (14) and with Eqs (7) and (14):

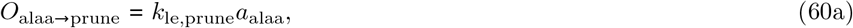

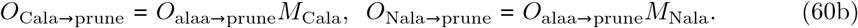

Units of the last two equations are kg aphid C, N stem^−1^ d^−1^.

##### Emigration

This flux, *O*_alaa→emi_ (aphid number stem^−1^ d^−1^) [Eq (61)], is equal to a standard temperature (20 °C) rate constant, *k*_alaa→emi20_, multiplied by a temperature-dependent function *f*(*T*_air_) for which the default parameter values are applied [Eq (17), Fig D.4]. Also, at low aphid densities [ρ_aph_, Eq (5)] so that the incoming alates do not immediately depart, emigration is switched off by multiplying by *f*_palaa→ala1_ [Eq (71), *cf*. Eq (26), Fig D.7B]. The combined rate parameter is (units: d^−1^)

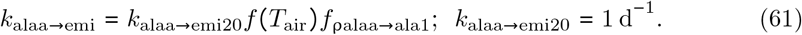

When the aphids are crowded, aphid density ρ_aph_ is large, *f*_palaa→ala1_ → 1 [Eq (71)] and emigration of adult alates occurs almost immediately. With the initial *t* = 0 values of *a*_ph_ = *a*_laa_ = 10 aphids per stem [Eqs Eq (64) and Eq (4)] and *A*_leaf,aph_ = 0.009 [Eq (5)], ρ_aph_ = 1111.1 [Eq (5)] and *f*_palaa→ala1_ = 0.552. Also at *t* = 0, *T*_air_ = 1.8 °C (Fig 3A), so that *f*(*T*_air_) = 0.019. Therefore [with Eq (7)]

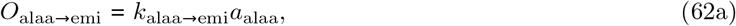

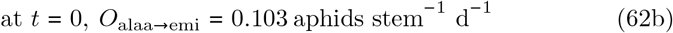

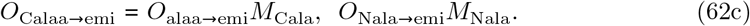

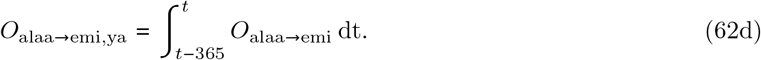

Alate adults (*a*_laa_, Fig 2) are the only morph that is able to emigrate. Units are aphids stem^−^ d^−^ and kg C, N stem^−1^ d^−^. Equation (62d) gives the yearly accumulation (ya) of emigrants (aphids stem^−1^ d^−1^). For the simulation shown in Fig 4, +0 °C which is chaotic, this also behaves chaotically. The total output of adult alate aphids (aphids stem^−1^ d^−1^) is obtained by adding the contributions [Eqs (55), (60) and (62); *cf*. Eq (15)]:

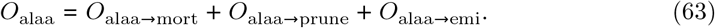

*O*_alaa_ is used in Eq (99) to calculate fractional outputs.

#### D.6.3 Differential equation: alate (winged) adult aphids

The differential equation (adult alates stem^−1^ d^−1^) and initial value for the state variable, a_laa_, are [with Eqs (57) and (63)]

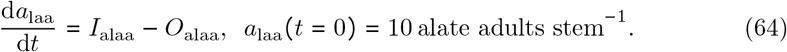

It is assumed that ten alate (winged) adult aphids fly in and colonize the stem at a specified time (usually *t* = 0 d).

### D.7 Alate aphid fecundity

The effect of temperature on alate fecundity is assumed to be the same as in Eq (18) for apterous aphid fecundity. The effect of nutrition on fecundity, manifest here through the N concentration in the phloem, *N*_phloem_ [Eq (2)], can be made specific — i.e. different for alate and apterous aphids, although here we assume apterous and alate adult aphids are equally fecund. The relevant equations are summarized below [*cf*. Eqs (19) to (24) for details]; results are given in Fig D.5.

The maximum volume intake rate of an alate adult, *v*_alaa, max_ (m^3^ aphid^−1^ d^−1^), places a limit on the maximum N intake from the phloem, giving a maximum phloem-volume-limited N intake [combining and modifying Eqs (19) and (20), but keeping the same parameter values]:

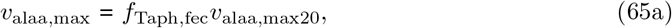

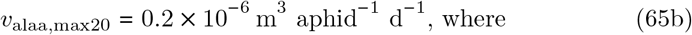

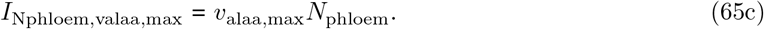

See Eq (18) for *f*_Taph,fec_. Units of the last equation are kg N aphid^−1^ d^−1^. The maximum amount of N which can be processed (*p*) is [*cf*. Eq(21)]

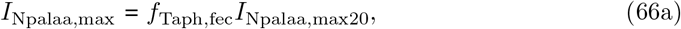

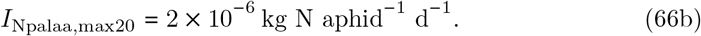

The N ingested per adult alate is the least of the volume-intake-limited [Eq (65)] or the processing-limited [Eq (66)] values (kg N aphid^−1^ d^−1^):

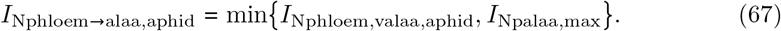

Dividing by *N*_phloem_ [Eq (2)], the actual volume of phloem sap ingested per alate adult aphid is (m^3^ phloem sap aphid^−1^) [*cf*. Eq (23)]

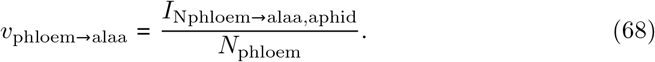

The C intake per aphid (kg C aphid^−1^ d^−1^) and then the N and C intakes per stem (kg N, C stem^−1^ d^−1^) are [*cf*. Eq (24)]:

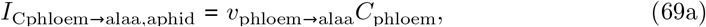

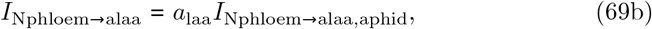

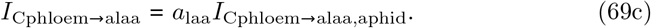

Here Eqs (68), (2), (64) and (67) are employed.

### D.8 Alate:apterous ratio in offspring from alate adults

The treatment above for the alate:apterous ratio in the offspring of apterous adults is followed exactly (Section D.3), but with the possibility of different parameters for the alates. Eqs (25)–(28) become

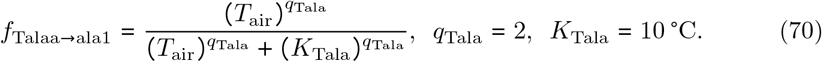

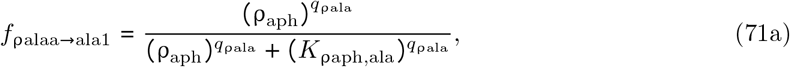

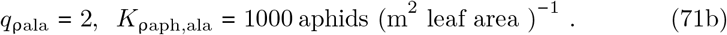

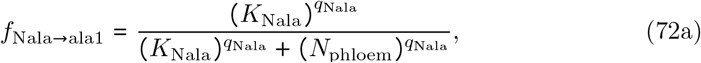

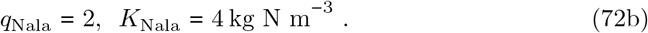

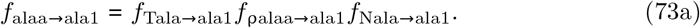

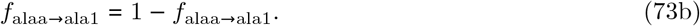

These quantities are drawn in Fig D.5 for the alate fractions of the progeny of apterous adults, where the parameterization is assumed the same as for alate adults.

#### D.8.1 Total fecundity and fractional fecundity of alate adults

Now that we have calculated the fractions of offspring from alate adults which are alate and which are apterous, *f*_alaa→ala1_ and *f*_alaa→apt1_ in Eq (73), we are able to calculate the total fecundity of alate adults, Oalaa→axx1 (1st instar aphids stem^−1^ d^−1^) [*cf*. Section D.3.4, Eq (29) for apterous adults].

The N ingested by alate adults [*I*_Nphloem→alaa_, (kg N stem^−1^ d^−1^), Eq (69)] is converted into alate and apterous first-instar nymphs, giving a total output of first-instar nymphs [*cf*. Eq (29)] and outputs and inputs to these pools with [*cf*. Eq (30)]

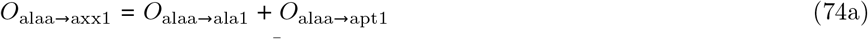

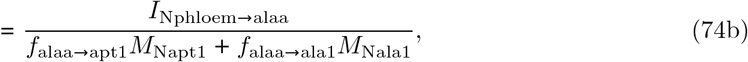

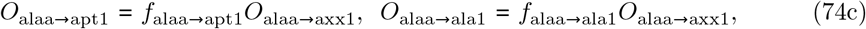

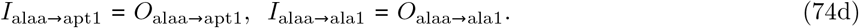

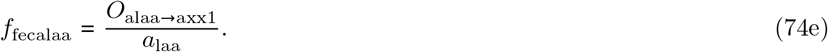

Here the total 1st instar progeny output from alate adults is *O*_alaa→axx1_ (aphids stem^−1^ d^−1^) and the total fractional fecundity of alate adults is *f*_fecalaa_ (d^−1^).

#### D.8.2 Associated N and C fluxes

The N fluxes associated with these fluxes of aphids are (kg N stem^−1^ d^−1^)

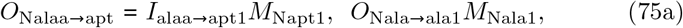

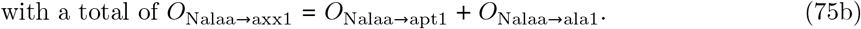

The carbon intake of the alate adults is *I*_Cphloem→alaa_ [Eq (69), (kg C stem^−1^ d^−1^)]. There are carbon fluxes (kg C stem^−1^ d^−1^) to respiration, offspring and the residual C flux goes to honeydew [*cf*. Eqs (32) to (34)]:

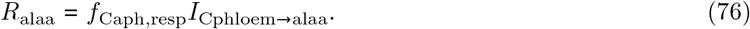

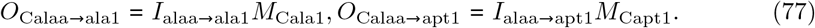

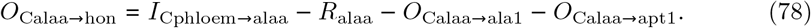

Here on the right side of Eqs (76) to (78) we have used Eqs (32), (69), (74) and (7). The honeydew output is input to the surface litter metabolic pool [see Eq (107) and above].

### D.9 Development rates of alate aphids

These are treated as in Eqs (35) to (37) (Section D.4) for apterous aphids, although development rates are slightly different for alate instars. We replace Eq (38) by

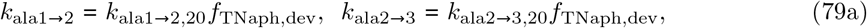

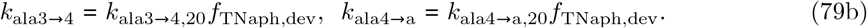

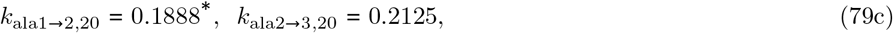

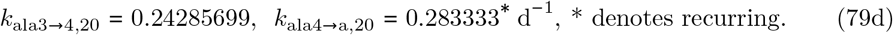

Here *f*_TNaph,dev_ is given by Eq (37). The lifetimes associated with these 20 °C rate constants are

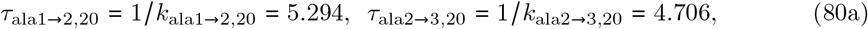

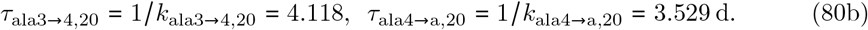

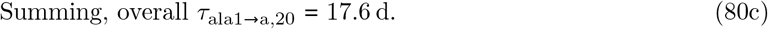

The overall value agrees reasonably with Dixon ([4], p. 109, their table 6.1). Alates develop 15% more slowly than apterous aphids [*loc. cit*., Dixon refers to Fisher [93] for data on the green spruce aphid]. See also Section D.4.1 on survival.

### D.10 Juvenile alate (winged) aphids

There are four instars, with state variables *a*_la1,…,4_ (aphids per stem) (Fig 2). The treatment is the same as above for the apterous instars (Section D.5) where the pools are treated together, defining first the inputs, then the outputs and last the differential equations. The equations are summarized below.

#### D.10.1 Inputs: juvenile alate (winged) aphids

Inputs, *I*_ala*j*_, *j* = 1,…, 4 (aphids stem^−1^ d^−1^), to these pools are [Fig 2, *cf*. Eq (41)]

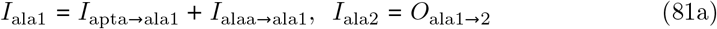

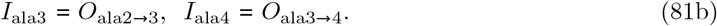

The inputs to the first pool are from the reproduction of both apterous and alate adults [Fig 2, Eqs (30) and (74). The inputs to the other three pools are equal to the developmental outputs of the preceding pool [Eq (82)]. It is assumed that transfer takes place without loss.

#### D.10.2 Outputs: juvenile alate (winged) aphids

The outputs, *O*_ala*j*_, *j* = 1,…, 4, arise from development, mortality and tree/foliage pruning. There is no loss of aphid numbers per stem to thinning when whole stems are removed.

##### Development

The developmental output fluxes from the four alate instars are [*cf*. Eq (42)] (aphids stem^−1^ d^−1^):

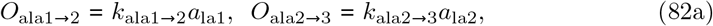

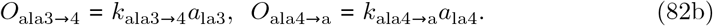

Rate constants are in Eq (79). State variables are defined in Eq (87). These outputs are assumed to occur without loss, providing the inputs (aphids stem^−1^ d^−1^) of *I*_ala2-4_ in Eq (81) and also a developmental input to alate adults [see Eq (57)] of

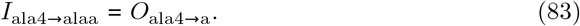

##### Mortality

These are [with Eqs (11) and (87); *cf*. Eq (44)] (aphids stem^−1^ d^−1^):

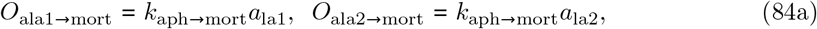

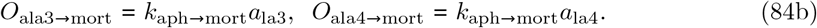

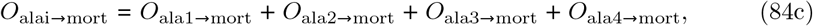

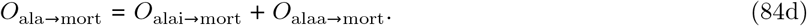

It is assumed here that the same mortality rate constant applies to alate and apterous instars. The last two of Eq (84) give the total alate instar and the total alate output to mortality.

##### Pruning

These are [with Eqs (14) and (87); *cf*. Eq (45)] (aphids stem^−1^ d^−1^):

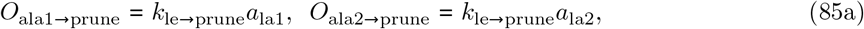

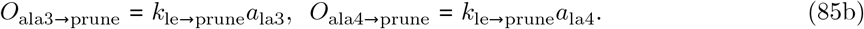

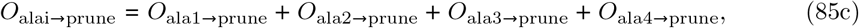

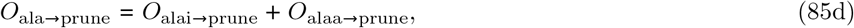

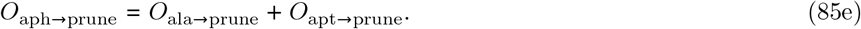

The last three of Eq (85) give first, the total alate instar output to pruning, *O*_alai→prune_; next, with the first of Eq (60), total output of alate aphids to pruning, *O*_ala→prune_; and last, the total aphid output to pruning, *O*_aph→prune_, using the last of Eq (45). See Section D.5.2 ‘pruning,’ for pruning of apterous aphids. The pruning rate constant *k*_le→prune_ = 0 d^−1^ in the aphid studies presented here [Eq (14)].

#### D.10.3 Differential equations: juvenile alate (winged) aphids

Total outputs for each of the four instar pools are [with Eqs (82), (84) and (85)] [*cf*. Eq (64)]

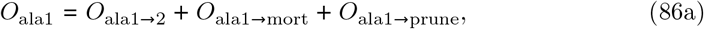

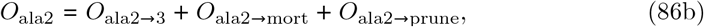

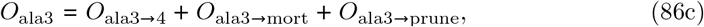

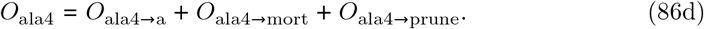

Combining Eqs (81) and (86), the differential equations and initial values for the four state variables are (aphids stem^−1^ d^−1^) [*cf*. Eq (47)]:

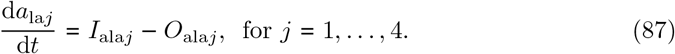

At *t* = 0 d, *a*_la1_ = *a*_la2_ = *a*_la3_ = *a*_la4_ = 0 aphids stem^−1^.

#### D.10.4 Nitrogen and carbon fluxes for the four alate instar pools

Assume the N required from phloem is the amount required to increase the N component of aphid mass entering the next pool. Therefore with Eq (82) [*cf*. Eq (48)]:

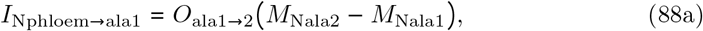

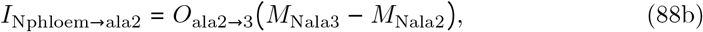

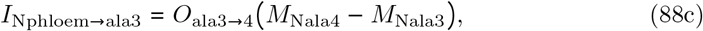

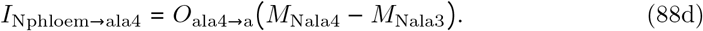

Units are: kg N stem^−1^ d^−1^. Aphid N contents are in Eq (7). Total N taken from the phloem by the four instars (alai) is

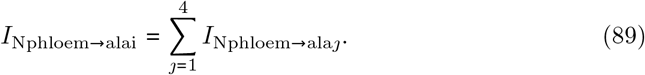

The C flux from the phloem accompanying this N flux is [Eq (3), *cf*. Eq (50)]:

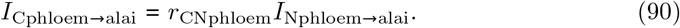

Respiration flux is (kg C stem^−1^ d^−1^)

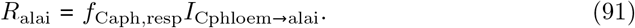

*f*_Caph,resp_ is given in Eq (32).

The increase in instar mass from aphid instar development also requires C from the phloem. The four input (*I*) components of this and the total phloem C output to supply these four inputs are [with Eqs (82) and (7); *cf*. Eq (52)] (kg C stem^−1^ d^−1^):

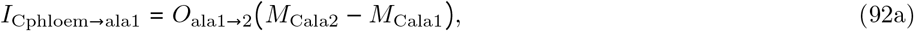

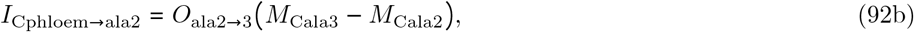

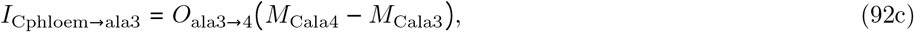

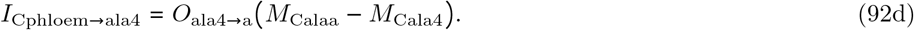

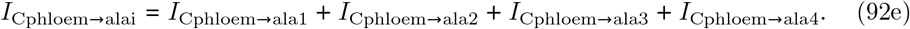

The excess C is excreted as honeydew (input to the surface litter metabolic pool, Eq (107), see Thornley [34], chapter 5, *cf*. Eq (53)]:

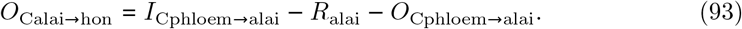

This makes use of Eqs (90)–(92).

Mortality gives rise to C and N outputs [Eqs (7) and (84), *cf*. Eq (54)] (kg C, N stem^−1^ d^−1^):

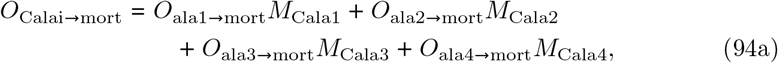

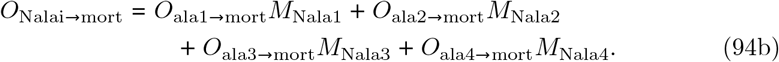

The C, N mortality fluxes above are captured in Eqs (108) and (118) and thence in Eqs (112) and (126) on the way to the surface litter cellulose and lignin pools [Eq (127)].

Pruning similarly gives rise to C and N outputs [with Eqs (85) and (7), *cf*. Eq (55)] (kg C, N stem^−1^ d^−1^):

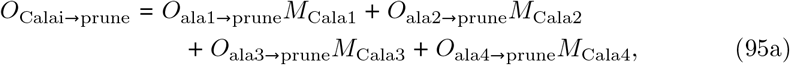

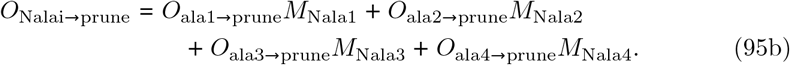

These fluxes are captured in Eqs (110) and (120) and thence in Eqs (112) and (112).

### D.11 Aphid mortality, emigration and fecundity

Here, we calculate total mortality, emigration and fecundity for the aphids.

#### D.11.1 Mortality

Aphid fluxes to mortality are summed to give a total of *O*_aph→mort_ (aphid number stem^−1^ d^−1^) of

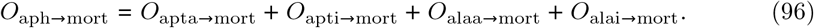

Here we have used Eqs (12), (44), (59) and (84). Because these fluxes vary greatly both diurnally and seasonally, it can be helpful to take yearly accumulations (*ya*) with

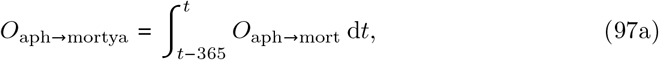

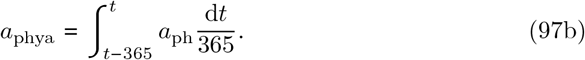

The additional subscript ya denotes yearly accumulation for rate variables such as *O*_aph→mort_ or a yearly average for non-rate variables such as *a*_ph_ · *O*_aph→mortya_ now has units of aphid number stem^−1^ y^−1^ · *a*_phya_ is a yearly average of *a*_ph_, with units of aphid number stem^−1^.

#### D.11.2 Emigration

The emigration flux is *O*_alaa→emi_ (aphids stem^−1^ d^−1^) [Eq (62)]. Total output from the alate adult pool is *O*_alaa_ [Eq (63)]. The mortality flux from this pool is *O*_alaa→mort_ [Eq (59)]. In Fig D.9 the fractional output fluxes to mortality and to emigration are shown. In the absence of pruning [Eq (63)] the two fractional variables defined in Eq (99) add up to unity. Emigration peaks in late summer, when the adult alate population is maximum (Fig D.9) and air temperature is also favourable (Fig 3A).

**Fig D.9.**
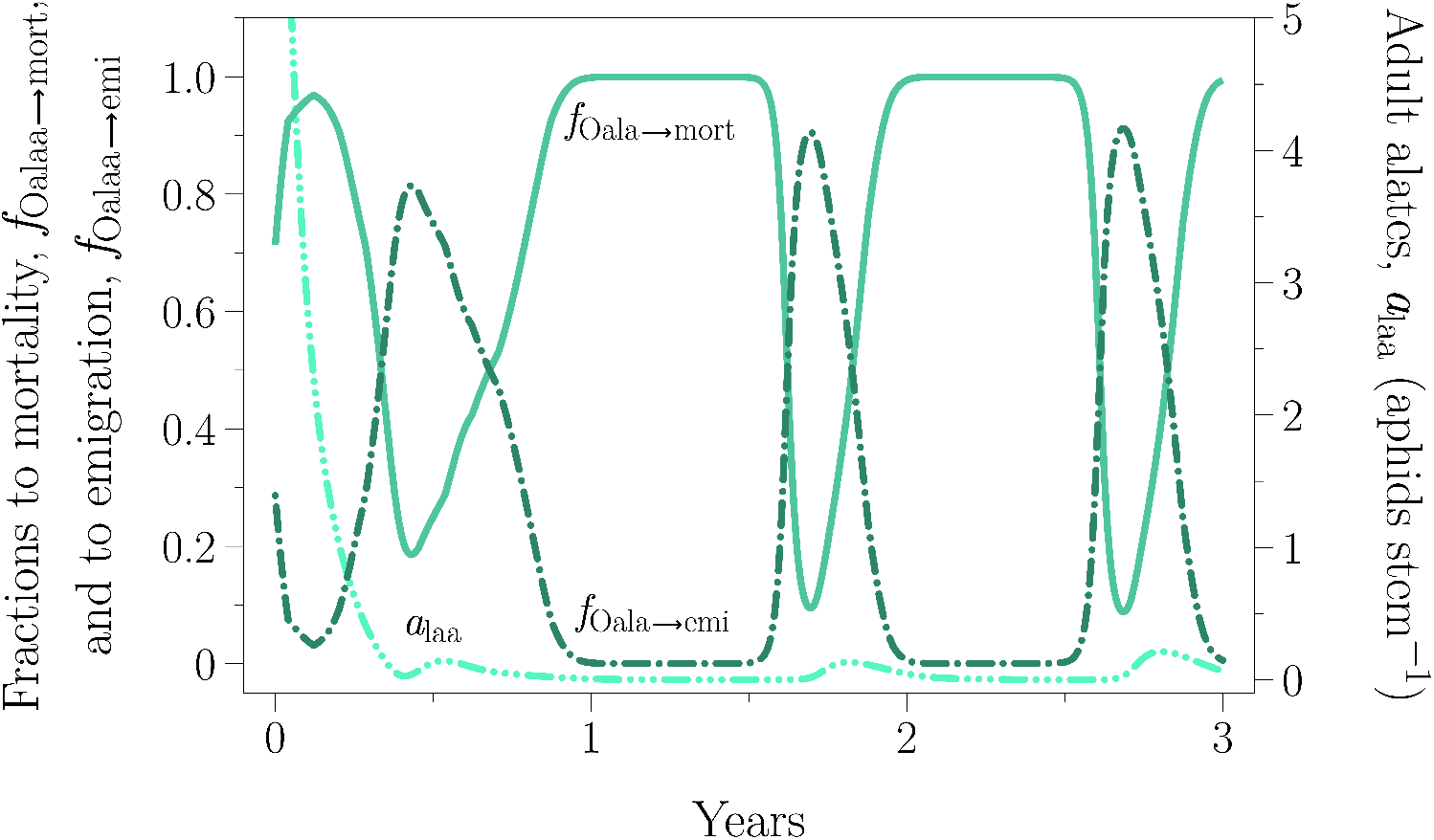
Fractional output fluxes. Fractional output fluxes of alate adults to mortality, *f*_Oalaa→mort_ (solid line) and to emigration, *f*_Oalaa→emi_ [Eq (99)] (dash-dot line) over three years post-infection [Eq (64)] at time *t* = 0 for a plantation in a northern Britain environment (Section 4.1). The state variable for alate adults a·alaa (Fig 2) is shown by the dashed-dot-dot-dot line [Eq (64)] with reference to the right side ordinate; initial value is 10 and is therefore not shown.

Because, as far as the local population is concerned, emigration is equivalent to mortality, we also calculate the flux to ‘mortality + emigration’, which includes the emigration flux from alate adults [*O*_alaa→emi_, Eq (62)]. Using an obvious notation, therefore

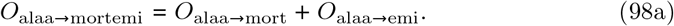

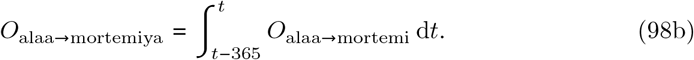

The terms Eq (98a) have units of aphids stem^−1^ d^−1^; in Eq (98b) we give the yearly accumulations (aphids stem^−1^ y^−1^). In fact, the emigration flux is mostly small compared to the total aphid mortality flux (Fig D.9). In Eq (98) and in the simulations shown in Fig D.9, mortality and emigration are the only fluxes from the alate adult pool [Eq (63)] as the pruning fluxes are zero [Eq (14)]. The (dimension-free) fractional outputs are

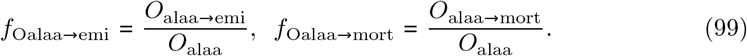

The fractional fate of alate adults is drawn in Fig D.9. Aphids in the alate adult pool, *a*_laa_, may either die or emigrate [Fig 2 and Eq (63). The fractions are calculated in Eq (99). After an initial transient the fraction emigrating, *f*_Oalaa→emi_ drops to a few percent and then peaks as the temperature rises [Fig 3A, Eq (17), Fig D.6, default temperature response]. Mortality rate is close to zero at these moderate temperatures [Eqs (11) and (9), Fig D.3]. Alate adult numbers per stem, *a*_laa_, increase each year as leaf area available to the aphids, *A*_leaf,aph_, increases [Eq (5)].

#### D.11.3 Fecundity

Total offspring production rate from apterous and alate adults (1st instar aphids stem^−1^ d^−1^) is

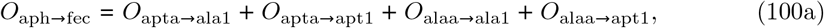

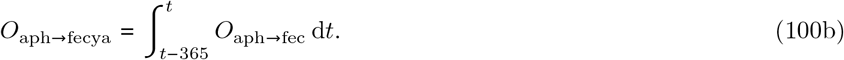

Here we utilized Eqs (30) and (74). Eq (100a) gives the instantaneous rate and Eq (100b) the yearly accumulation (1st instar aphids stem^−1^ y^−1^).

#### D.11.4 Total aphid number per stem, *a*_ph_

The ten state variable differential-equation components of *a*_ph_ (Fig 2) of Eq 4, namely Eqs (64), (87), (16) and (47) are added to give d*a*_ph_/d*t*:

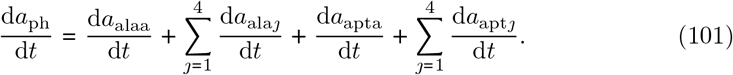

Putting immigration, *I*_imm→alaa_ = 0 [Eq (56)] and noting that alate adults (*a*_laa_) are the only morph able to emigrate, therefore

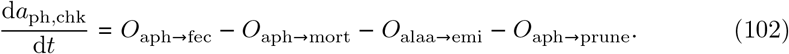

Units are aphids stem^−1^ d^−1^. Here we make use of Eqs (100), (96), (62) and (85). The equality of the values of Eqs (101) and (102) gives a useful check on the accuracy of the programming.

### D.12 Aphid sub-model — C and N balances

In this section the inputs and outputs of the whole aphid subsystem are summarized. This has two purposes. First, it is a useful check that the model has been consistently formulated mathematically and is programmed correctly. Second, in long-term ecosystem studies, quite small fluxes are important, as they can lead to degradation, or otherwise, of the ecosystem. These are now made explicit.

#### D.12.1 Carbon inputs: phloem, immigration

There are two C inputs to the aphid sub-model, from the phloem and from possible immigration of alates. The C input from the phloem is [Eqs (90), (50), (69) and (24)]

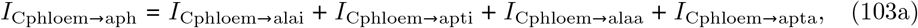

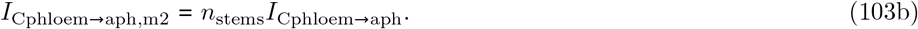

The first equation is in units of kg C stem^−1^ d^−1^, the second, kg C (m^−2^ ground)^−1^ d^−1^. *n*_stems_ is the stem density (number of stems m^−2^), a state variable of the efm [Eq (1)]. Immigration gives C fluxes of [Eqs (7) and (56)]

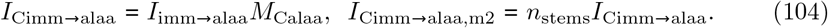

The first equation is in units of kg C stem^−1^ d^−1^, the second, kg C (m^2^ ground)^−1^ d^−1^. Total C inputs to the aphid sub-model are [Eqs (103) and (104)]

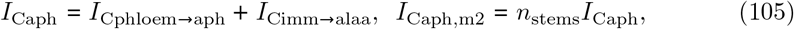

with units of kg C stem^−1^ d^−1^ and kg C (m^2^ ground)^−1^ d^−1^.

#### D.12.2 Carbon outputs: respiration, honeydew, mortality, emigration, pruning, thinning

There are outputs via respiration, honeydew, mortality, emigration, pruning and thinning.

Aphid respiration which produces CO_2_ gives C fluxes of [Eqs (76), (91), (32), (51)]

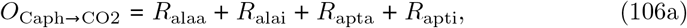

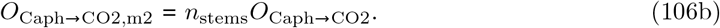

Units are kg C stem^−1^ d^−1^ and kg C (m^2^ ground)^−1^ d^−1^, respectively.

Honeydew production [Eqs (78), (93), (34), (53)] gives an output, *O*_Caph→hon,m2_, which is input without loss to the surface litter metabolic pool [Eq (12)] ([34], figure 5.1, *C*_surf-li,met_):

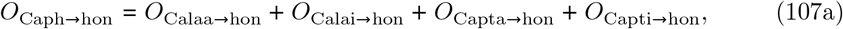

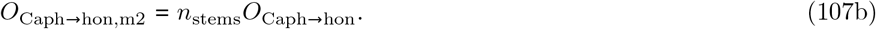

Units are kg C stem^−1^ d^−1^ (Eq (107a)) and kg C (m^2^ ground)^−1^ d^−1^ (Eq (107b)). Mortality of aphids gives rise to C fluxes of [Eqs (59), (94), (12) and (54)]

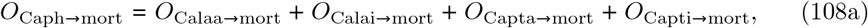

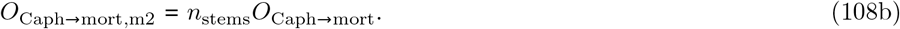

Units are as in Eq (107). This flux is divided equally between the surface litter cellulose and lignin pools [Eq (126) and Eq (127); figure 5.1 in [34]].

Emigration of adult alates gives rise to a output C flux (kg C stem^−1^ d^−1^) of *O*_Calaa→emi_ [Eq (62)] and a C flux per unit area ground of

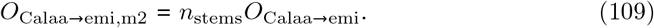

Pruning the trees gives an output C flux of [Eqs (60), (95), (14), (55)]

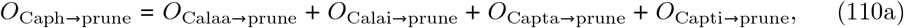

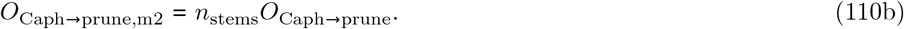

Again, units are kg C stem^−1^ d^−1^ and kg C (m^2^ ground)^−1^ d^−1^. As with the mortality fluxes of Eq (108), it is assumed that pruning produces a flux of dead aphids into the surface litter cellulose and lignin pools [Eqs (126) and (127); figure 5.1 in [34]].

Thinning, which is applied in forest plantation management, removes whole stems (Section 4.2). It does not affect aphid numbers per stem. However, aphids on removed stems are assumed to join the surface litter, together with the foliage on the removed stems (some of the branches and stems may be taken away as timber products). The rate of stem loss per unit ground area is specified by the management regime in the forest model (Section 4.2). It is *O*_nstems→th_ [Eq (1), stems m^−2^ d^−1^]. This quantity is zero most of the time, but it is non-zero on the days when the plantation is thinned by an amount specified. Multiplying by *C*_aph_ [Eq (6), kg aphid C per stem], this leads to an aphid C output flux per unit ground area of (kg C m^−2^ d^−1^)

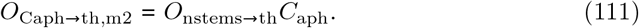

This output is input to the soil surface litter inputs in Eq (126).

Total aphid C outputs are [with Eqs (106)–(108), (110), (62) and (111)]

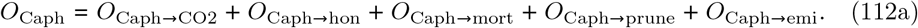

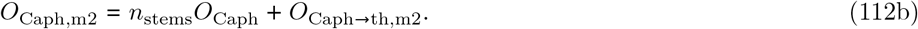

The first equation here has units of kg C stem^−1^ d^−1^; the second units of kg C (m^2^ ground)^−1^ d^−1^.

#### D.12.3 Carbon balance

A consistency check is to calculate the rate of change of aphid C two ways. First by summing the differential Eqs (64), (87)), (16) and (47), with Eq (7) for the first equation and Eqs (1) and (6) for the second equation:

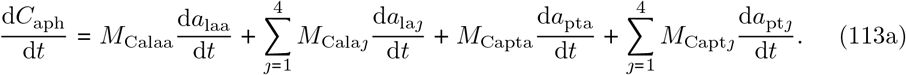

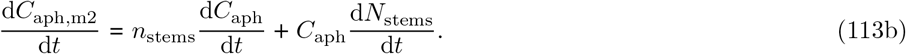

These were compared (successfully) with the same (check = chk) quantities calculated with [Eqs (105) and (112)]

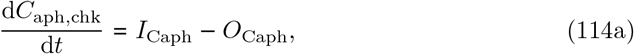

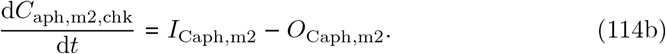

#### D.12.4 Nitrogen inputs: phloem, immigration

The procedure from Eqs (103)–(114) for C is repeated for N. There are the same two N inputs [*cf*. Eqs (103)–(105)], from the phloem and via immigration.

From the phloem, therefore, with Eqs (89), (49), (69), (24) and (1) for *n*_stems_:

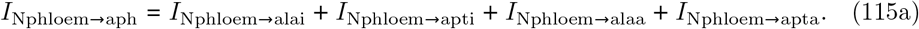

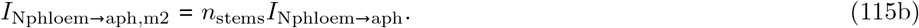

Units are kg N stem^−1^ d^−1^ and kg N (m^2^ ground)^−1^ d^−1^.

Immigration gives N fluxes of [Eqs (7) and (56)]:

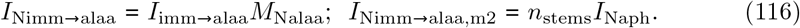

Units as in Eq (115).

Total N inputs to the aphid sub-model are [with Eqs (115), (116) and (1)]

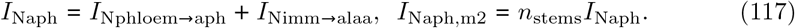

Units are kg N stem^−1^ d^−1^ and kg N (m^2^ ground)^−1^ d^−1^.

#### D.12.5 Nitrogen outputs: mortality, emigration, pruning, thinning

There are outputs to mortality, emigration, pruning and thinning.

First, mortality of aphids gives rise to N fluxes of [Eqs (59), (94), (13), (54) and (1)]

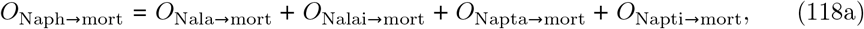

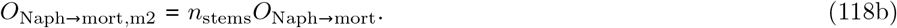

Units are kg N stem^−1^ d^−1^ and kg N (m^2^ ground)^−1^ d^−1^. This dead aphid N flux is divided between the surface litter metabolic N, cellulose and lignin pools (Eqs (126)–(128); figure 5.1 in [34]).

Next emigration of adult alates gives rise to an N flux per stem of *O*_Nalaa→emi_ [Eq (62)] and an N flux per unit area ground of [for *n*_stems_, see Eq (1)]:

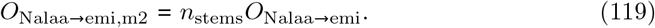

Pruning the trees gives an output N flux of [Eqs (60), (95), (14) and (55)]

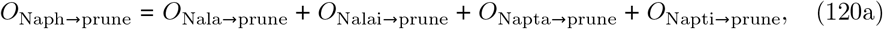

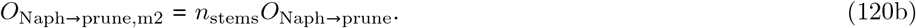

Units are kg N stem^−1^ d^−1^ and kg N (m^2^ ground)^−1^ d^−1^.

Thinning gives rise to an aphid N output flux per unit ground area of [Eq (6), see discussion before Eq (111)] (kg N m^−2^ d^−1^)

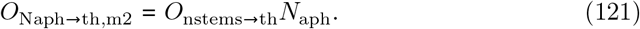

Total aphid N outputs are [with Eqs (118), (120), (62), (1) and (121)]

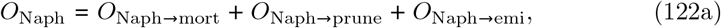

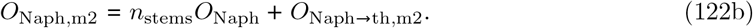

Units are kg N stem^−1^ d^−1^ and kg N (m^2^ ground)^−1^ d^−1^. These outputs are used in Eq (124) for consistency checking and in Eq (126) for calculating inputs to the surface litter pools.

#### D.12.6 N balance

The same consistency check is applied to N as for C [Eqs (113) and (114)].

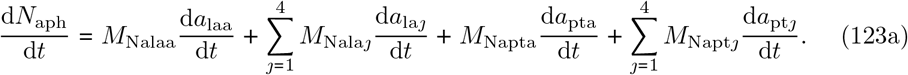

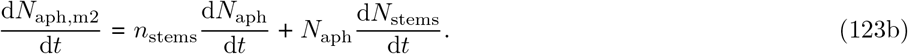

Units of these two equations are kg N stem^−1^ d^−1^ and kg N (m^2^ ground)^−1^ d^−1^. For the first of these sum the differential Eqs (64), (87), (16) and (47) and apply Eq (7); for the second use Eqs (1) and (6). These were compared (successfully) with those calculated with Eqs (117) and (122).

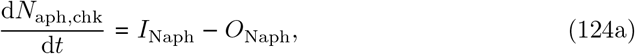

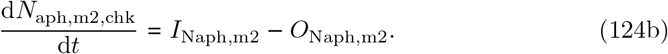

### D.13 Soil sub-model inputs from: mortality, pruning and thinning

The C input to the surface litter metabolic C pool ([34], figure 5.1) from honeydew is [Eq (107)] (kg C m^−2^ d^−1^):

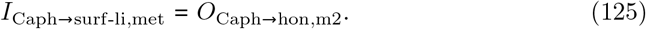

The C inputs to the cellulose and lignin pools are calculated by first totalling the C and N dead aphid fluxes from mortality [Eqs (108) and (118)], pruning [Eqs (110) and (120)] and thinning [Eqs (111) and (121)] (kg C, N m^−2^ d^−1^):

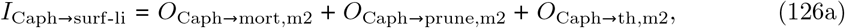

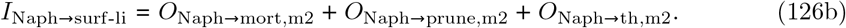

The C flux is partitioned equally between the cellulose and lignin pools:

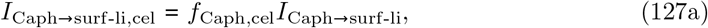

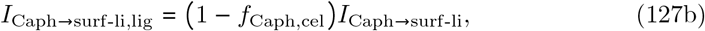

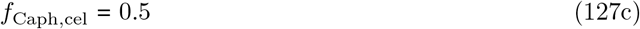

*f*_Caph,cel_ is the fraction of C in dead aphids assigned to the surface litter cellulose pool, the remainder entering the lignin pool.

The N flux is handled rather differently, because of the assumption that the cellulose and lignin litter pools have a constant C:N ratio ([34], p. 75; p. 85, equation 5.1c). The N flux required to satisfy this fixed C:N ratio is subtracted from the total dead aphid N flux and the remainder is put into the surface litter metabolic pool. Therefore, with Eqs (127) and (126):

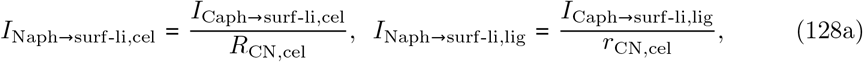

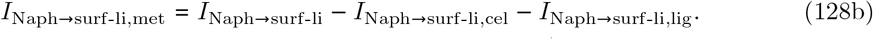

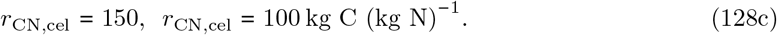

The cellulose and lignin C:N ratios are 150 and 100 kg C (kg N)^−1^.

### D.14 Aphid C and N at the end of a rotation

At the end of a rotation, the plantation is clear-felled. It is assumed that the aphid C and N are similar to foliage structural C and N and they are added to the surface litter structural C and N pools on this basis, before the next rotation is begun. See figure 5.1 and equations 5.1b,c of Thornley ([34], p. 76, p. 85).

When initializing for the next rotation, the initial C and N contents of the aphids (*C*_aph_, *N*_aph_ at *t* = 0, expressed per unit ground area) are subtracted from the surface litter metabolic C and N pools, so that small amounts of C and N are not added at the beginning of every rotation when infestation via alate immigration occurs. However this is not important in the present studies as the initial values for the soil and litter sub-model are obtained by multiple rotations in the absence of aphids.

## Appendix E Summary of symbols used in the model

**Tab E.1.**
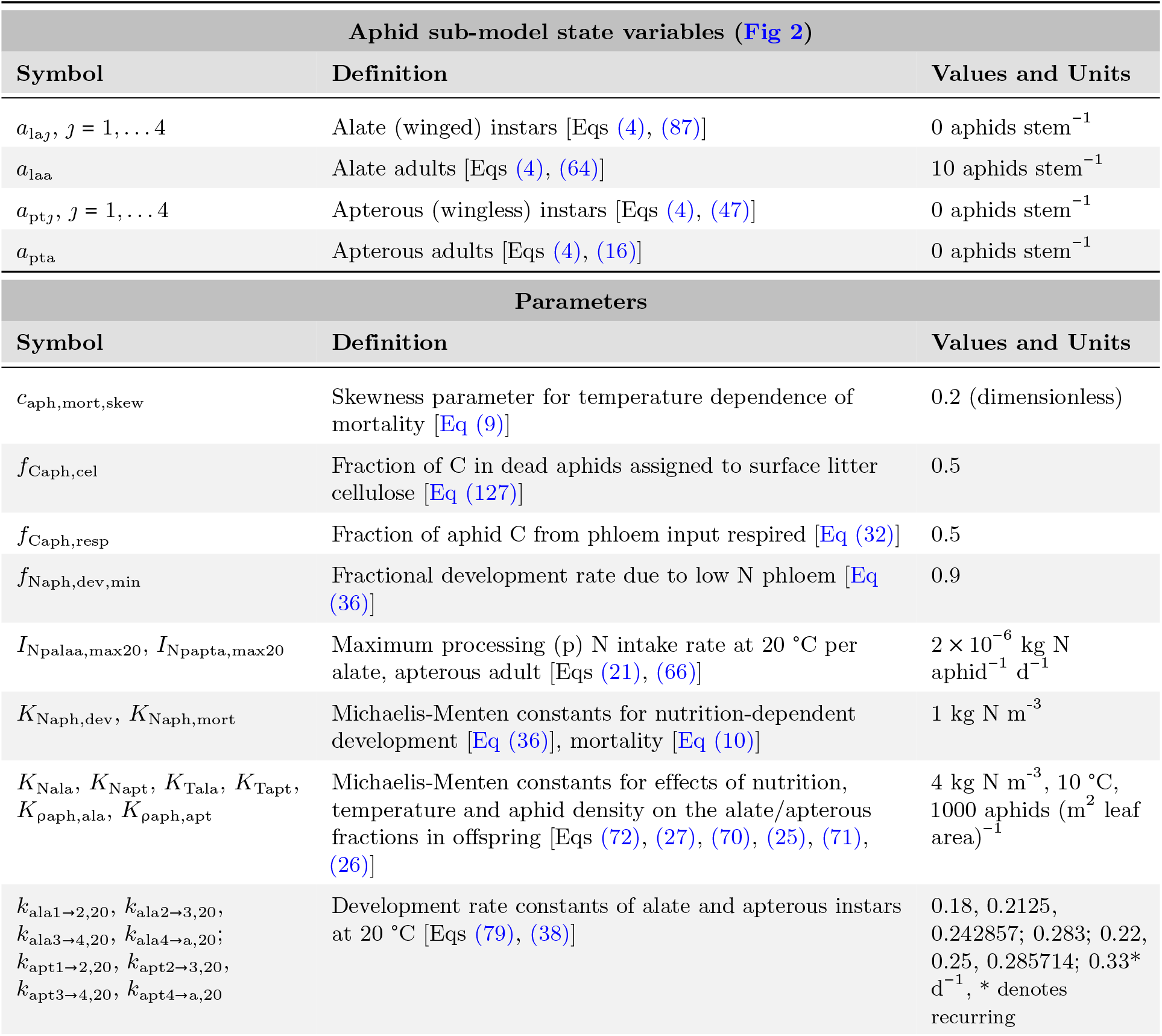
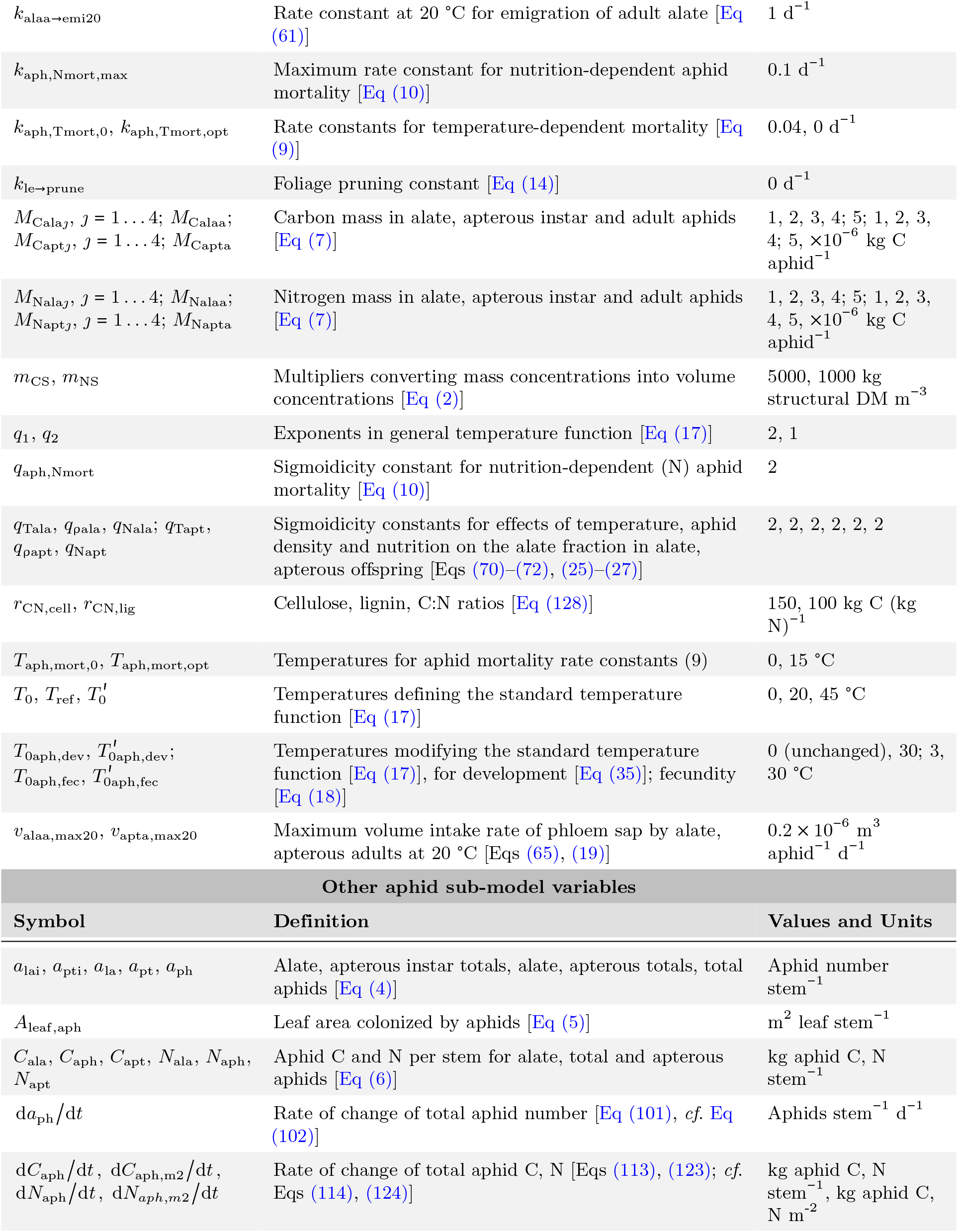
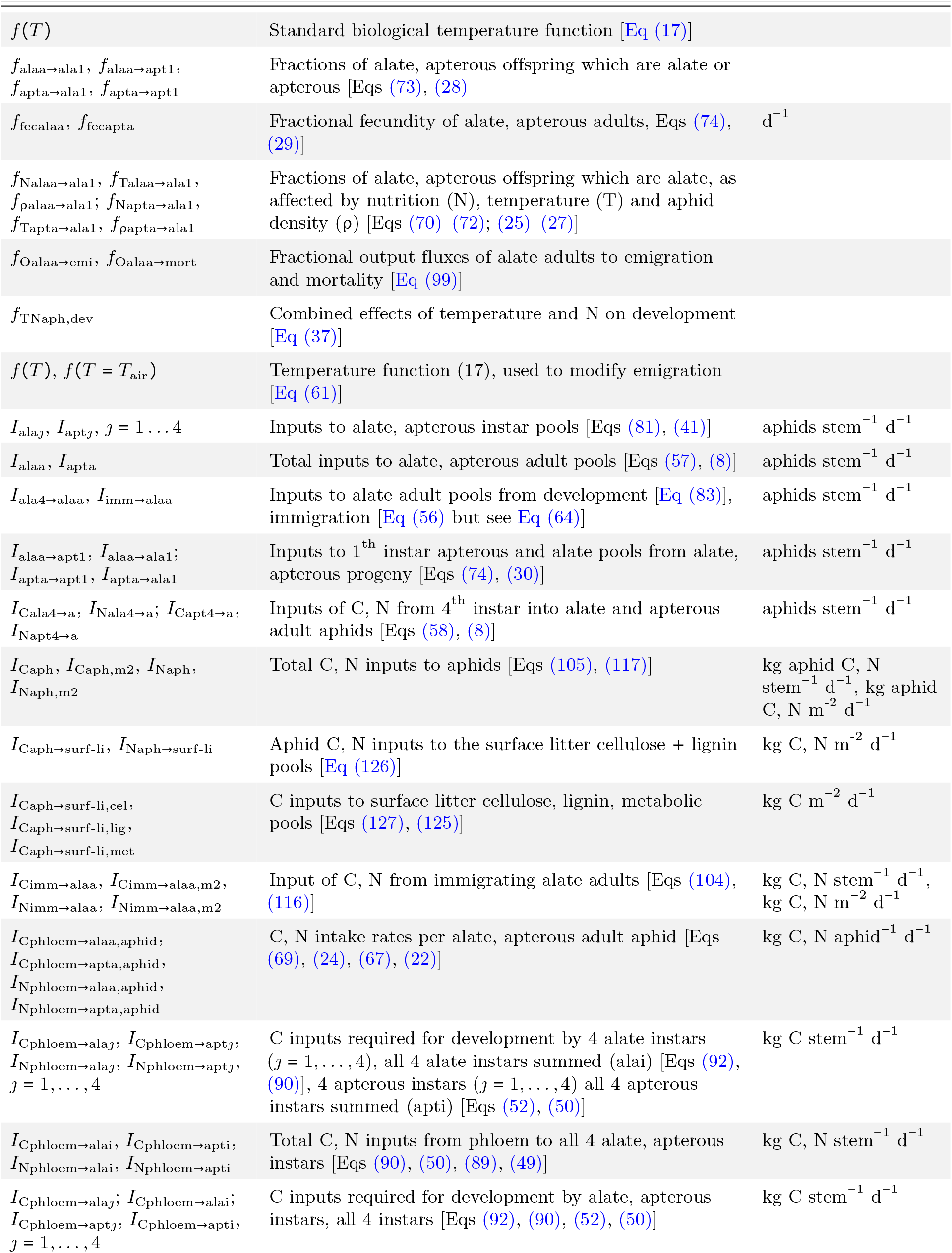
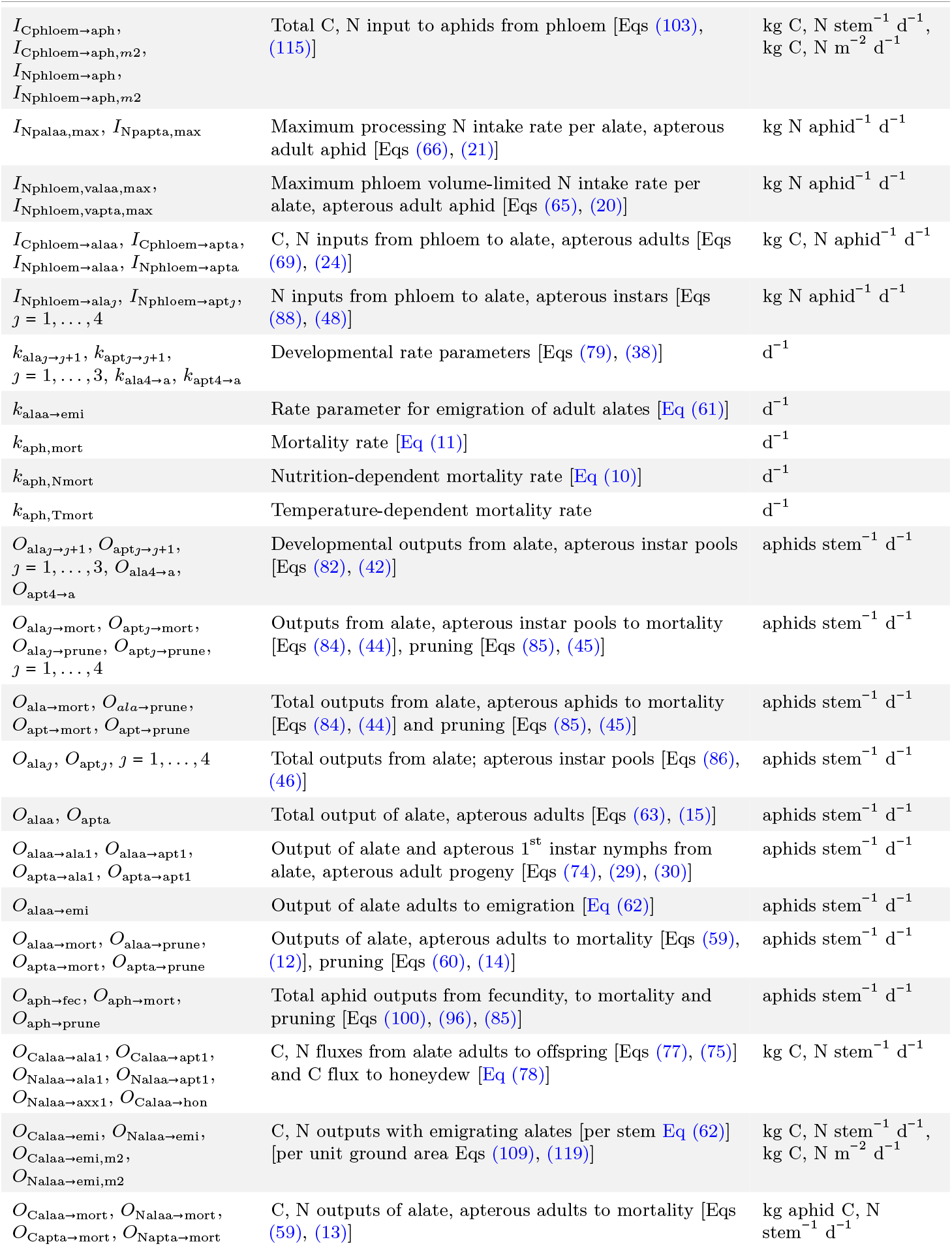
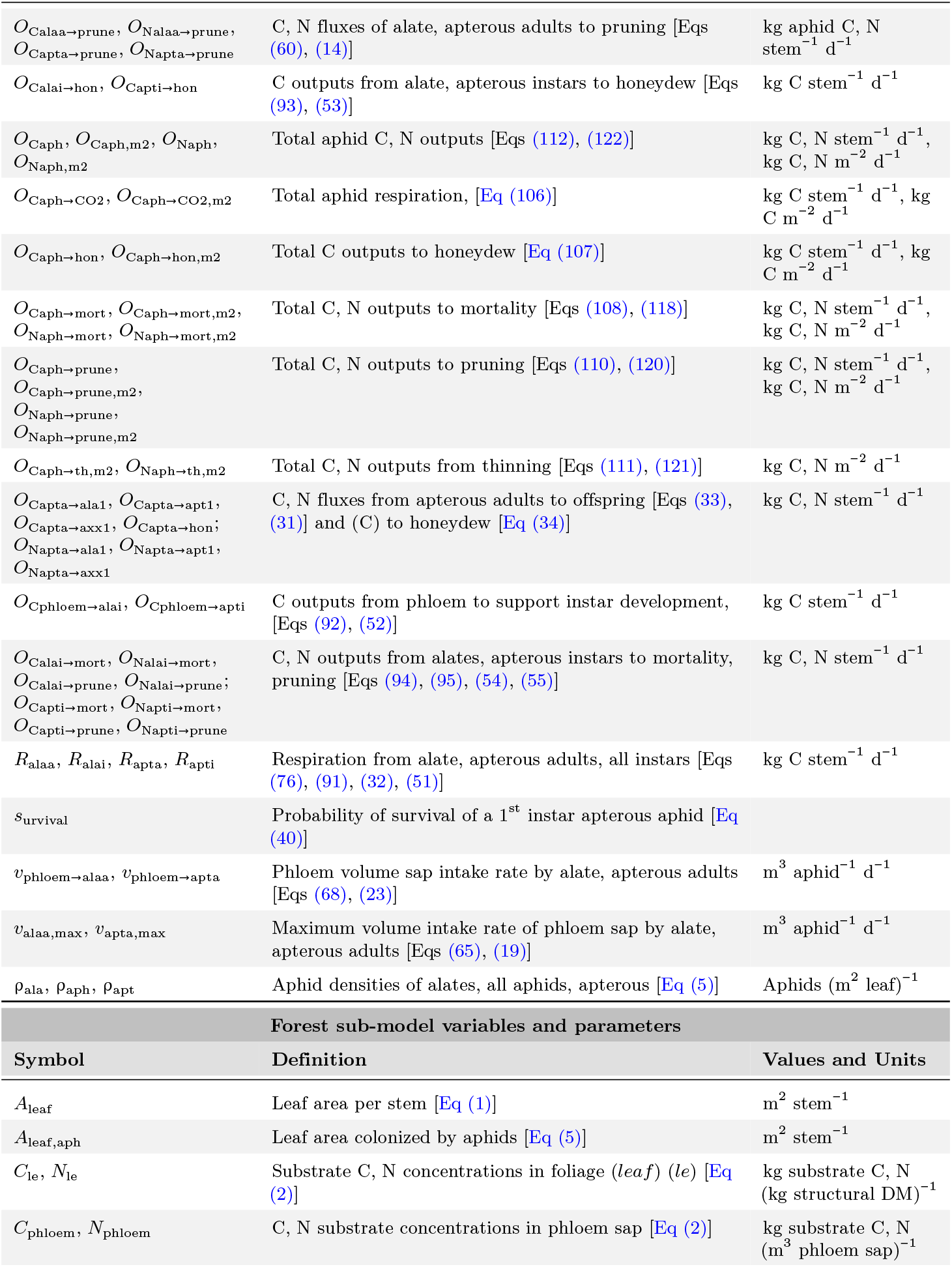
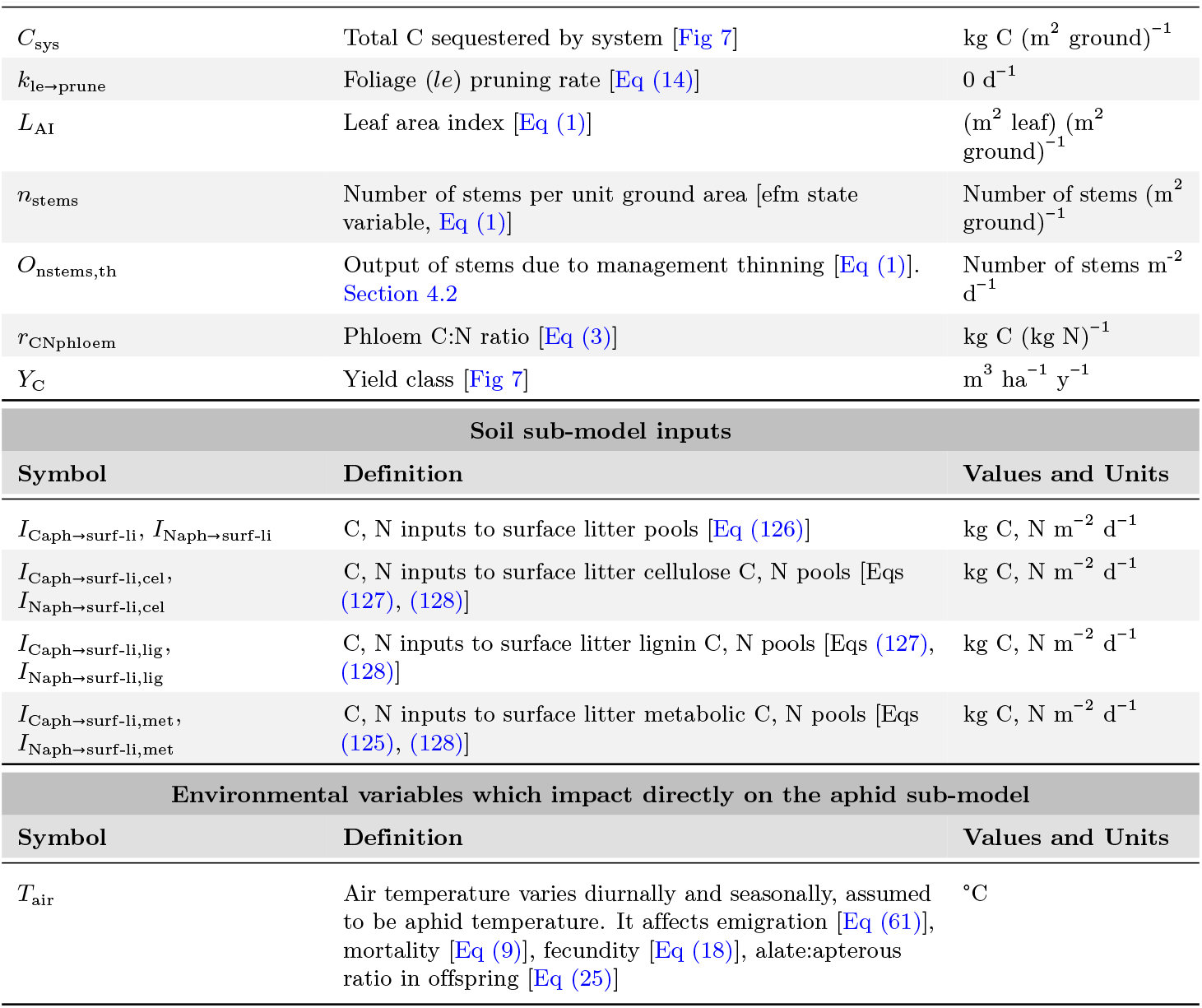
Symbols and definitions. Notation and abbreviations: equation numbers of significant occurrence are given. DM denotes dry matter, C denotes carbon, and N denotes nitrogen.

## Footnotes

1 see https://www.jmp.com/support/help/en/16.0/index.shtml#page/jmp/statistical-details-for-spectral-density.shtml

